# CART-GPT: A T Cell-Informed AI Linguistic Framework for Interpreting Neurotoxicity and Therapeutic Outcomes in CAR-T Therapy

**DOI:** 10.1101/2025.08.08.669387

**Authors:** Tiantian Mao, Xiaojian Shao, Wei Guo, Zhiwu Jiang, Rui Jing, Xin Li, Yiran Zhu, Tony Jin, Tao Ma, Yong Lu, Guangxu Jin

**Affiliations:** Houston Methodist Cancer Center/Weill Cornell Medicine, Houston, TX 77030, USA; Digital Technologies Research Center, National Research Council Canada, Ottawa, ON, K1A 0R6, Canada; Department of Biochemistry, Microbiology and Immunology, Ottawa Institute of Systems Biology, University of Ottawa, Ottawa, ON, K1H 8M5, Canada; Atkins Academic & Technology High School, Winston Salem, NC 27101, USA; Department of Internal Medicine-Gerontology and Geriatric Medicine, Wake Forest University School of Medicine, Winston-Salem, North Carolina, USA; Comprehensive Cancer Center, Atrium Health Wake Forest Baptist, Winston-Salem, NC 27157, USA; Department of Cancer Biology, Wake Forest University School of Medicine, Winston-Salem, NC 27157, USA

**Author notes:** These authors contribute to this work equally (Tiantian Mao, Xiaojian Shao, Wei Guo, Zhiwu Jiang, Rui Jing, Xin Li).

**Keywords:** AI models in immunotherapy, large language model–based prediction, ICANS risk prediction, CAR-T outcome prediction

## Abstract

Chimeric antigen receptor (CAR) T cell therapy holds transformative potential for hematologic malignancies, yet predicting patient-specific treatment efficacy and neurotoxicity remains a major clinical challenge due to the complex and heterogeneous nature of the infused CAR-T cell populations. Here, we introduce CART-GPT, a transformer-based model fine-tuned on a curated atlas of 1.12 million CAR-T single-cell RNA-seq profiles annotated with clinical outcomes. CART-GPT is the first AI model developed for CAR-T therapy that predicts both treatment response and the risk of immune effector cell-associated neurotoxicity syndrome (ICANS), achieving state-of-the-art performance (AUC ~0.8) and marking a significant advance in the field. The model provides interpretable insights, revealing that neither therapeutic efficacy nor neurotoxicity is driven by individual cell types alone, but by the combined influence of discrete, distinct subsets across diverse T cell states and transcriptional programs. A novel cell aggregation strategy links single-cell predictions to patient-level metrics, enhancing both accuracy and biological relevance. As a contribution to this ever-evolving field, we also release a comprehensive, annotated single-cell CAR-T atlas as a community resource to facilitate future research in immunotherapy. These advances demonstrate the potential of foundation models in single-cell biology to inform precision CAR-T treatment planning and facilitate the rational design of next-generation cell therapies.

## Introduction

Chimeric antigen receptor (CAR) T cell therapy has transformed the treatment landscape for hematologic malignancies, particularly B cell cancers^1–3^. Despite the remarkable advances over the past few decades, broader clinical implementation of CAR-T therapy remains hindered by substantial challenges, most notably, heterogeneous treatment responses and severe immune-related toxicities^4,5^. For instance, in non-Hodgkin lymphoma, up to 40% of patients deemed eligible for CAR T cell therapy ultimately receive treatment but are later identified as non-responders^6,7^. Meanwhile, immune effector cell–associated neurotoxicity syndrome (ICANS) is a particularly concerning adverse effect, with high-grade ICANS (grade 3–4) observed in approximately 25–32% of treated patients^8–10^. Alarmingly, some patients not only fail to benefit from therapy but also develop life-threatening neurotoxicity. Consequently, the ability to understand and predict treatment efficacy and toxicity early—ideally prior to or at the time of infusion—would greatly enhance patient stratification and inform more effective, personalized treatment strategies.

Single-cell RNA sequencing (scRNA-seq) has enabled unprecedented profiling of cellular heterogeneity at infusion and post-infusion stages, offering new opportunities to decipher the transcriptional programs and cellular compositions that underlie patient-specific treatment outcomes^11,12^. Previous studies have identified correlations between certain T cell subsets—such as memory-like CD8^+^ or central memory CD4^+^ T cells—and treatment outcomes, as well as associations between T cell activation or exhaustion markers and ICANS^13–16^. However, these findings fall short of capturing the complex, individualized immune dynamics that govern CAR-T therapy outcomes. Despite the promise of scRNA-seq, leveraging it for clinical prediction remains challenging due to its high dimensionality, technical noise, and biological variability. A key barrier lies in translating cell-level insights into reliable patient-level predictions. Current approaches often treat cells in isolation or fail to account for the combinatorial behavior of diverse cell states within a patient’s immune ecosystem. Critically, there is a lack of robust, scalable frameworks that can integrate single-cell data to stratify patients by expected therapeutic benefit or toxicity risk. This underscores the urgent need for models that can bridge cell- and patient-level information, resolve complex immune signatures, and ultimately guide individualized treatment decisions in CAR-T therapy, including intensified neurological monitoring or early intervention with corticosteroids or IL-6 blockade (e.g., tocilizumab) ^4,5^.

Recent advances in artificial intelligence (AI), particularly transformer-based large language models (LLMs) trained on large-scale single-cell RNA sequencing (scRNA-seq) data, offer a promising framework for modeling complex cellular behaviors^17–19^. However, applying these broadly trained AI models to predict clinical responses of CAR-T cell therapy remains an unmet goal. This is due in part to the difficulty of adapting general-purpose architectures to CAR-T-specific tasks, which requires carefully curated data and task-specific optimization. Bridging this gap by fine-tuning LLMs on CAR-T scRNA-seq data and linking single-cell states to patient-level responses could unlock new opportunities to decode the cellular “language” of CAR-T therapy through AI-powered modeling.

Here, we introduce CART-GPT, an integrated AI toolbox built upon a fine-tuned transformer foundation model, specifically designed to interpret the cellular language of CAR-T therapy. Leveraging a new comprehensive CAR-T cell atlas of over 1.12 million cells annotated with treatment response and ICANS outcomes, CART-GPT comprises three functional modules: (1) TcellGPT for high-resolution T cell subtype annotation, (2) CART-GPT-response for predicting treatment efficacy, and (3) CART-GPT-ICANS for stratifying ICANS risk. Our findings suggest that both therapeutic efficacy and ICANS are not determined by individual cell types alone, but rather by the intricate latent interplay of diverse T cell states, forming an interpretable immune “language” that CART-GPT is uniquely equipped to decode. This work holds the potential to transform CAR-T therapy by enabling precision-guided interventions and ultimately enhancing both patient safety and clinical outcomes.

## Results

### CAR-T-Cell Atlas

To build a foundation for modeling CAR-T cell states and predicting clinical outcomes, we first utilized a large reference T cell atlas consisting of approximately 216,000 T cells across 17 human tissues, sourced from publicly available single-cell RNA-seq datasets^20^ (**Figure 1a, Supplementary Table 1**). This atlas includes a wide range of immune-relevant tissues such as blood, bone marrow, and kidney, and captures diverse T cell subtypes, including CD4^+^ helper T cells, CD8^+^ alpha-beta memory T cells, and gamma-delta T cells. In addition, we manually curated a CAR-T cell atlas from the infusion products (IP) of patients in published studies, comprising more than 1.12 million cells from 255 individuals **(Extended Data Figure 1)**. This CAR-T atlas includes 782,558 cells annotated with treatment response labels and 343,061 cells annotated with ICANS severity labels. By integrating both foundational and clinically annotated datasets, we established a robust resource to enable fine-grained modeling of CAR-T cell phenotypes and facilitate the development of predictive tools for clinical outcomes.

**Figure 1.**
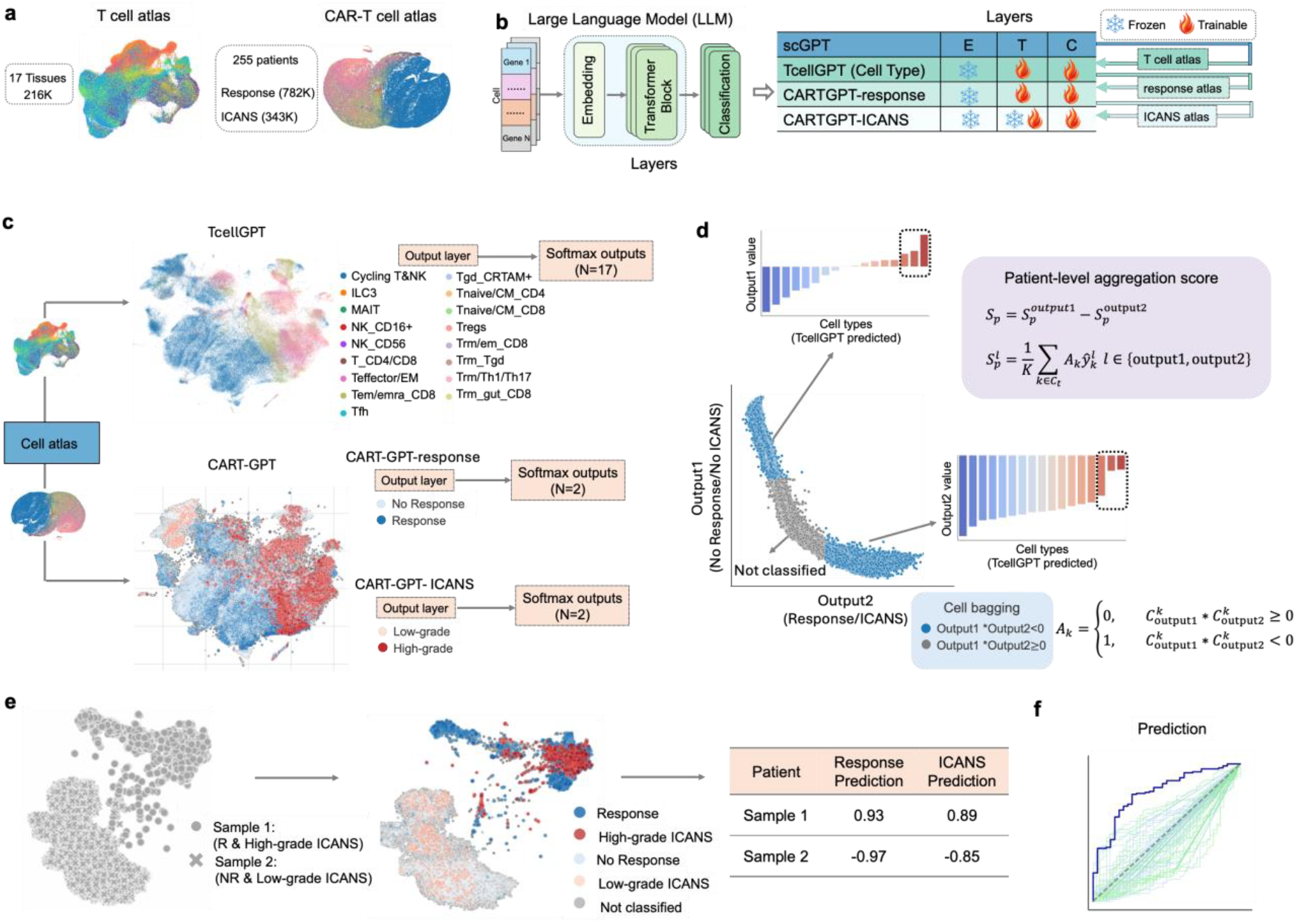
Schematic of the CART-GPT all-in-one AI toolbox. a, Construction of the reference atlases for model fine-tuning. A large T cell atlas was assembled from public databases, and a CAR-T cell atlas was manually curated from published studies. b, Architecture of CART-GPT. The model builds upon the pretrained scGPT by freezing its embedding layers and fine-tuning the remaining components to create TcellGPT, a general-purpose model for T cell annotation. TcellGPT was then further trained on CAR-T data to generate CART-GPT-response for predicting therapy response. Thus, TcellGPT was subsequently fine-tuned into CART-GPT-ICANS. c. CART-GPT tools and their outputs in the output layers. d, Illustration of the “cell bagging” strategy. Cells are scored based on classification confidence derived from the model’s classification layers. Ambiguous cells with uncertain or conflicting predictions are excluded, and only high-confidence cells are aggregated to compute robust patient-level prediction scores. e, Example visualizations of model-predicted distributions for ICANS and response. Left: Sample 1: good response; high ICANS; Sample 2: no response; low ICANS. Right: model-predicted cells; blue: response; light blue: no-response; red: high ICANS; light red: low-ICANS. f, Overview of the CART-GPT clinical outcome prediction.

### CART-GPT Overview: All-in-One AI toolbox for CAR-T cell therapy

CART-GPT is a fine-tuned extension of a transformer-based foundation model (scGPT)^19^, trained on a large-scale single-cell atlas that includes both natural T cells and CAR-T data, as described above. It provides an all-in-one toolbox for T cell subtype annotation, CAR-T cell therapy response prediction, and ICANS risk evaluation (**Figure 1b**). Leveraging transfer learning, CART-GPT provides three specialized tools: (1) TcellGPT, which decodes CAR-T cell states within the broader context of natural T cell biology, (2) CART-GPT-response, which predicts patient-specific treatment efficacy, and (3) CART-GPT-ICANS, which predicts the likelihood of neurotoxicity (**Figure 1c, Methods**). TcellGPT serves as the foundation for CART-GPT-response and CART-GPT-ICANS training. Separately, CART-GPT-response and CART-GPT-ICANS implement a novel “cell bagging” strategy to bridge single-cell predictions with patient-level outcomes (**Figure 1d**). Rather than treating all cells equally, this approach intelligently aggregates cell-level outputs by identifying and prioritizing high-confidence, informative subpopulations—those most predictive of response or ICANS. By focusing on these aggregated cells, the model generates patient-level metrics that emphasize biologically relevant signals while minimizing noise (**Figure 1e**). Ultimately, we evaluate the performance of the CART-GPT tools in predicting treatment response and ICANS severity with robust patient-level resolution (**Figure 1f**).

### TcellGPT: T cell subtype annotation

To enable fine-grained classification of T cell states, we fine-tuned the pretrained scGPT model using a curated T cell atlas (**Supplementary Table 1**), resulting in TcellGPT, an auto-annotation tool for T cell subtype prediction. TcellGPT incorporates 12 transformer layers, 512-dimensional cell embeddings, and outputs 17 subtype classes. Evaluation against manually curated annotations showed that TcellGPT achieved high prediction accuracy across all subtypes, with overall accuracy exceeding 90% (**Extended Data Figure 2a**). The predicted subtype distribution closely mirrored manual annotations in both rank order and relative abundance—for instance, Tnaive/CM_CD4 was the most abundant subtype (over 15%), while ILC3 was the least (less than 1%), a pattern faithfully recapitulated by TcellGPT (**Extended Data Figure 2b–c, Supplementary Table 2**).

To assess model applicability, we compared TcellGPT to CellTypist^21,22^, a widely used and informative cell type annotation tool designed for general-purpose classification. While both tools accurately predict T cell populations, they operate at different levels of resolution. For example, TcellGPT distinguishes fine-grained subsets such as T follicular helper (Tfh) cells from broader helper T cell populations, whereas CellTypist provides more general T cell annotations. For shared populations, such as Cycling T&NK, MAIT, and ILC cells, both tools produced comparable distributions across these cell types (**Extended Data Figure 2e–g**). Given that CellTypist is optimized for broad cell annotation, some differences in subtype resolution may arise when compared to manually curated T cell-specific labels (**Extended Data Figure 2d**).

Overall, TcellGPT offers a complementary approach that aligns closely with manual annotations, making it especially well-suited for high-resolution characterization of T cell subsets.

### CART-GPT-response: predicting CAR-T cell therapy response

CART-GPT-response fully leverages TcellGPT and the cell bagging framework to predict CAR-T cell therapy response. In this approach, TcellGPT first annotates T cell subtypes and its model architecture, as well as its pretrained weights—specifically for the transformer backbone—are transferred to initialize CART-GPT-response. Building on this, the architecture for CART-GPT-response preserved the core components of TcellGPT, consisting of 512-dimensional input embedding layers followed by 12 stacked transformer blocks, while modifying the classification output layer to include two output nodes instead of the original 17 (**Figure 2a**). During training, the input embedding layers are retained without modification, while the transformer and classification layers are fine-tuned. This transfer learning strategy from TcellGPT to CART-GPT-response preserves the model’s ability to represent T cell states while enabling adaptation to the new task of predicting treatment response. To train and validate the model, we assembled a dataset comprising over 780,000 cells from 199 patients, using a patient-level five-fold cross-validation (**Figure 2b, Extended Data Figure 1b-c**).

**Figure 2.**
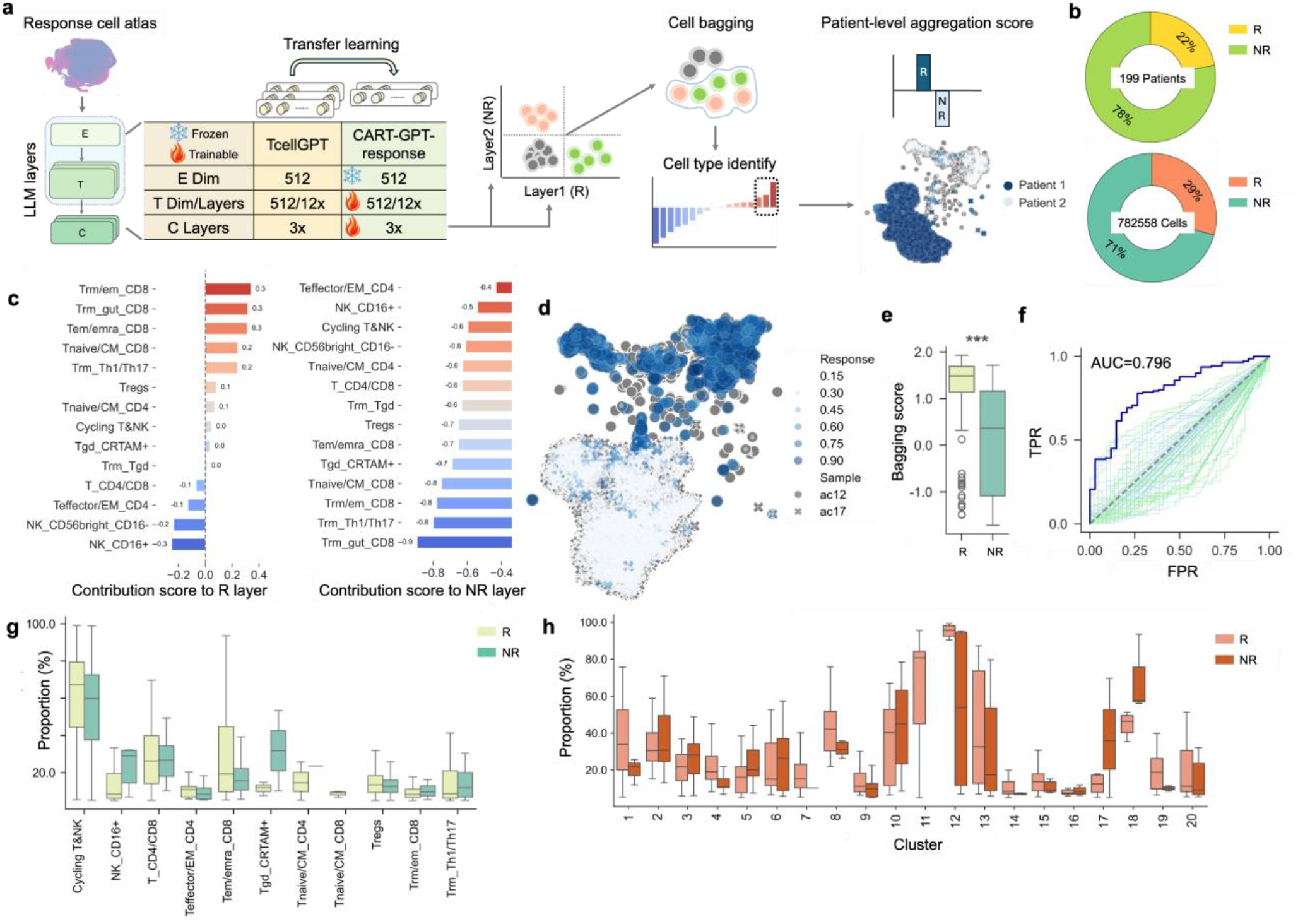
CART-GPT-response model for predicting CAR-T cell therapy response. a, Schematic of the CART-GPT-response model architecture. Starting from TcellGPT, the model was fine-tuned using CAR-T response-labeled single-cell data and adapted to output two classification nodes: one for responders (R group) and one for non-responders (NR group). b, Dataset summary. The response dataset includes 782,558 cells from 199 patients, each annotated with clinical response outcomes and used for model training and evaluation. c, Cell-type contributions to response prediction. Bar plots display T cell subtypes with the highest contribution scores to the R classification node (left) and NR classification node (right). The X-axis represents contribution score; the Y-axis indicates cell types predicted by TcellGPT. d, Patient-specific classification signatures. Visualization of cells from two representative patients—ac12 (responder) and ac17 (non-responder)—colored by per-cell contribution scores. Larger, darker points represent higher contributions; ○ represent patient ac12 and cross markers × represent patient ac17. e, Patient-level bagging scores for outcome classification. Boxplots show aggregated bagging scores for responders and non-responders. The X-axis denotes the two groups: light yellow green indicates responders and teal indicates non-responders; the Y-axis shows bagging score values. ^***^ *P* value < 10^−6^, Mann-Whitney U test. f, Model performance evaluation. The dark blue curve displays the ROC curve for CART-GPT-response with an AUC of 0.796. Light green lines indicate the ROC curves of cell types; light blue lines indicate the ROC curves of Leiden clusters. g, T cell subtype distribution between responders and non-responders. Boxplots compare the proportion of each TcellGPT-predicted cell type across patients. The X-axis shows the proportion of each cell type within individual patients; the Y-axis lists predicted T cell subtypes. Light yellow green corresponds to responders; teal corresponds to non-responders. Each bar in the plot represents the proportion of this cell type within an individual, with longer bars indicating substantial heterogeneity in cell abundance within the group. h, Unsupervised cluster composition by response group. Cells are grouped by UMAP-derived unsupervised clusters; the X-axis represents cluster identities; the Y-axis shows the proportion of each cluster within each patient. Light salmon pink bars represent responders; dark orange-brown bars represent non-responders. Each bar in the plot represents the proportion of this cell type within an individual, with longer bars indicating substantial heterogeneity in cell abundance within the group.

Following the removal of ambiguous cells using the cell bagging strategy, previously underrepresented cell types—such as Trm_Th1/Th17 cells—emerged as significant contributors to response prediction, highlighting their potential relevance despite their low abundance in the infusion product (**Figure 2c**). We further explored cell contributions at the individual patient level. Notably, patient ac12 (a responder) and patient ac17 (a non-responder) exhibited markedly different contribution patterns across the model’s classification layers, with distinct underlying cell-type distributions shaping their predicted outcomes (**Figure 2d, Extended Data Figure 3**). In the classification outputs, ac12 showed stronger contributions to the responder-associated class, reflecting a higher proportion of informative cell subpopulations relevant to therapeutic response. In contrast, ac17 exhibited limited contribution to this class and a more homogeneous cell composition. The more diverse contributing cells and stronger classification signal in ac12 enabled the model to accurately identify this patient as a responder, whereas the limited predictive signal in ac17 introduced greater ambiguity. Using CART-GPT-response, patients could be effectively stratified into responders and non-responders based on bagging scores, which reflect the confidence-weighted contribution of high-quality multiple cell subsets (**Figure 2e**). This approach enabled robust prediction of clinical response at the patient level, achieving a testing ROC-AUC of approximately 0.8, underscoring the model’s potential utility for outcome prediction in CAR-T cell therapy (**Figure 2f**).

While previous studies have linked specific cell types or gene expression patterns to clinical outcomes—such as memory-like CD8^+^ T cells^22,23^, CD4^+^ central memory T cells^23^, or the CD4^+^:CD8^+^ cluster ratio^24^—these associations, though informative, were insufficient to capture the underlying immune dynamics. For instance, cell types such as Trm/em_CD8, Teffector/EM_CD4, and Th1/Th17 were identified by CART-GPT-response as strong contributors to clinical outcome. However, their overall frequencies did not differ significantly between responders and non-responders (AUC = 0.29-0.68, ARI = –0.05-0.37; **Figure 2f–g**; **Supplementary Table 3**). In contrast, Tem/emra_CD8 cells exhibited high variability among responders. This observation highlights that predictive value is not necessarily tied to the average abundance of specific cell types across groups, but rather to their context-dependent roles within individual patients, underscoring a disconnect between cell type abundance and functional relevance. Moreover, classifying responders and non-responders based solely on unsupervised clustering, such as Leiden clustering^25^, proved challenging, as the overall cell distributions appeared similar between the two. Substantial intra-group variability highlights the limitations of using cluster abundance as a discriminative feature for classifying clinical response groups (AUC = 0.31-0.67, ARI = –0.12-0.25; **Figure 2f**,**h**; **Supplementary Table 3**). Together, these results highlight the ability of CART-GPT-response to capture nuanced, context-specific semantic features of CAR-T cells, going beyond surface-level annotations to deliver robust and interpretable predictions of therapeutic response.

### CART-GPT-ICANS: predicting neurotoxicity associated with CAR-T cell therapy

Building on the success of CART-GPT-response in predicting treatment efficacy, we further fine-tuned the model to develop CART-GPT-ICANS, aimed at predicting the risk of ICANS. This model was initialized from CART-GPT-response and further trained on our curated ICANS cell atlas, with architectural modifications including a reduction of the input embedding dimension from 512 to 64, a four-layer transformer block with 64-dimensional hidden states, and a three-layer classifier with a two-node output layer (**Figure 3a**). Similar to the response prediction strategy, CART-GPT-ICANS leverages cell bagging to aggregate high-confidence, ICANS-relevant cells into a patient-level aggregation score, enabling stratification of patients into high-risk (grade 3–4) and low-risk (grade 0–2) ICANS categories. The ICANS cell atlas used for training comprised over 343,000 cells from 56 patients, with 46% classified as high-grade (grade 3–4) and 54% as low-grade (grade 0–2) (**Figure 3b**).

**Figure 3.**
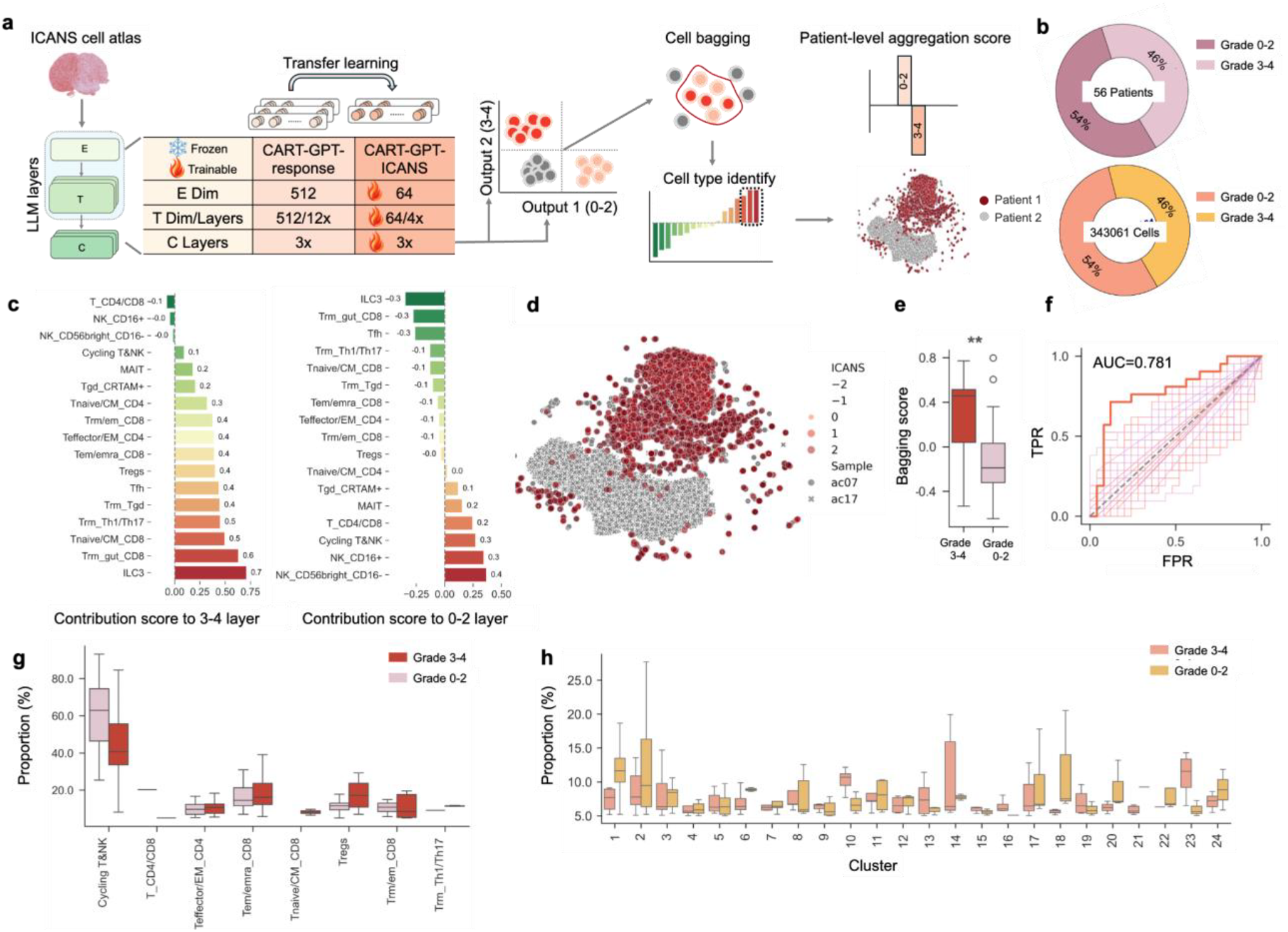
CART-GPT-ICANS model for predicting CAR-T cell therapy ICANS. a, Schematic of the CART-GPT-ICANS model architecture. Starting from CART-GPT-response, the model was further fine-tuned using ICANS-labeled single-cell data and adapted to output two classification nodes: one for high-risk patients (Grade 3–4 node) and one for low-risk patients (Grade 0–2 node). b, Dataset summary. The ICANS dataset includes 343,061 cells from 56 patients, each annotated with a clinical neurotoxicity grade. c, Cell-type contributions to ICANS prediction. Bar plots show T cell subtypes with the highest contribution scores to the Grade 3–4 group (left) and Grade 0–2 group (right). The X-axis represents contribution scores; the Y-axis indicates cell types predicted by TcellGPT. d, Patient-specific classification signatures. Visualization of cells from two representative patients—ac07 (Grade 3–4) and ac17 (Grade 0–2)—colored by per-cell contribution scores. Larger, darker points indicate stronger contributions. Blue shades represent contributions to response classification; red shades represent contributions to ICANS classification. ○ indicate ac07; × indicate ac17. e, Patient-level bagging scores for ICANS classification. Boxplots show aggregated bagging scores stratified by ICANS grade. The X-axis shows the two groups: dark red for Grade 3–4 (high risk) and lavender for Grade 0–2 (low risk); the Y-axis shows the bagging score. ^**^ *P* value < 10^−4^, Mann-Whitney U test. f, Model performance evaluation. The dark blue line shows the ROC curve of CART-GPT-ICANS for binary ICANS risk prediction, with an AUC of 0.781. Light purple lines indicate the ROC curves of cell types, muted red-pink lines indicate the ROC curves of Leiden clusters. g, T cell subtype distribution between high- and low-risk patients. Boxplots compare the relative proportions of TcellGPT-predicted cell types across patients. The X-axis shows the proportion of each cell type within individual patients; the Y-axis lists the predicted T cell subtypes. Dark red corresponds to Grade 3–4 patients; lavender corresponds to Grade 0–2 patients. Each bar in the plot represents the proportion of this cell type within an individual, with longer bars indicating substantial heterogeneity in cell abundance within the group. h, Unsupervised cluster composition by response group. Cells are grouped by UMAP-derived unsupervised clusters; the X-axis represents cluster identities; the Y-axis shows the proportion of each cluster within each patient. Light salmon pink bars represent high-risk patients; goldenrod bars represent low-risk patients. Each bar in the plot represents the proportion of this cell type within an individual, with longer bars indicating substantial heterogeneity in cell abundance within the group.

Model interpretation of the classification outputs revealed that specific cell types—including Trm/Th1/Th17 and Tem/emra_CD8—were strongly associated with high ICANS risk (**Figure 3c**), suggesting a potential immunological link between these subpopulations and neurotoxicity outcomes. To further illustrate the interpretability of CART-GPT-ICANS, we examined two representative patients: ac07 (high ICANS risk) and ac17 (low ICANS risk). The model revealed clear differences in cell-type contributions between these two patients (**Figure 3d, Extended Data Figure 4**). Notably, even within the high-risk patient ac07, not all cells contributed equally to the ICANS classification. Instead, the model selectively prioritized context-specific, high-impact cells, highlighting CART-GPT-ICANS’s ability to move beyond bulk abundance and focus on functionally relevant cell states that drive neurotoxicity risk. By aggregating these key cells through bagging, CART-GPT-ICANS successfully stratified patients into high- and low-risk groups (**Figure 3e**). The model achieved strong predictive performance, with a testing ROC-AUC of 0.78 by a 3-fold cross-validation (**Figure 3f**), demonstrating its potential utility for anticipating ICANS risk in CAR-T therapy recipients.

CART-GPT-ICANS revealed that cell-type abundance alone is insufficient to predict ICANS risk (AUC = 0.29-0.70, ARI = –0.01-0.01; **Figure 3f-g**; **Supplementary Table 4**), as overall frequencies and Leiden-based clustering failed to distinguish high (grade 3–4) and low-risk (grade 0–2) groups due to substantial intra-group heterogeneity (AUC = 0.30-0.68, ARI = 0-0.01; **Figure 3f**,**h**; **Supplementary Table 4**). These findings highlight the necessity of context-aware models like CART-GPT-ICANS, which incorporate predictive signals from individual cells to infer patient-level risk.

### CART-GPT decodes subtype-specific signals distinctly for response and neurotoxicity

To further assess the interpretability and robustness of CART-GPT tools across clinical endpoints, we evaluated a dataset in which both CAR-T therapy response and ICANS severity labels were available for the same cohort of patients^26^. This dual-labeled dataset included 51 patients with response outcomes (responder or non-responder) and 46 of these with ICANS severity annotations (grade 0-2 or grade 3-4), enabling joint evaluation of CART-GPT-response and CART-GPT-ICANS. Using the model trained from cross-validation fold 1, CART-GPT-response accurately predicted outcomes in 46 out of 51 patients (90.2%), misclassifying only two responders and three non-responders (**Figure 4a**). Interestingly, T cell subtype distributions varied considerably among responders, and in some cases resembled those of non-responders (**Figure 4b**). Nonetheless, the model effectively captured subtle immunological differences sufficient to correctly classify these patients. Using the model trained from cross-validation fold 1, CART-GPT-ICANS achieved 84.8% accuracy, correctly stratifying 39 out of 46 patients by ICANS severity, with three high-risk patients misclassified as low risk, and four low-risk patients misclassified as high risk (**Figure 4c**). Compared to response classification, ICANS prediction proved more challenging due to the similarity in cell type distributions between high- and low-risk groups (**Figure 4d**), emphasizing the need for fine-grained modeling.

**Figure 4.**
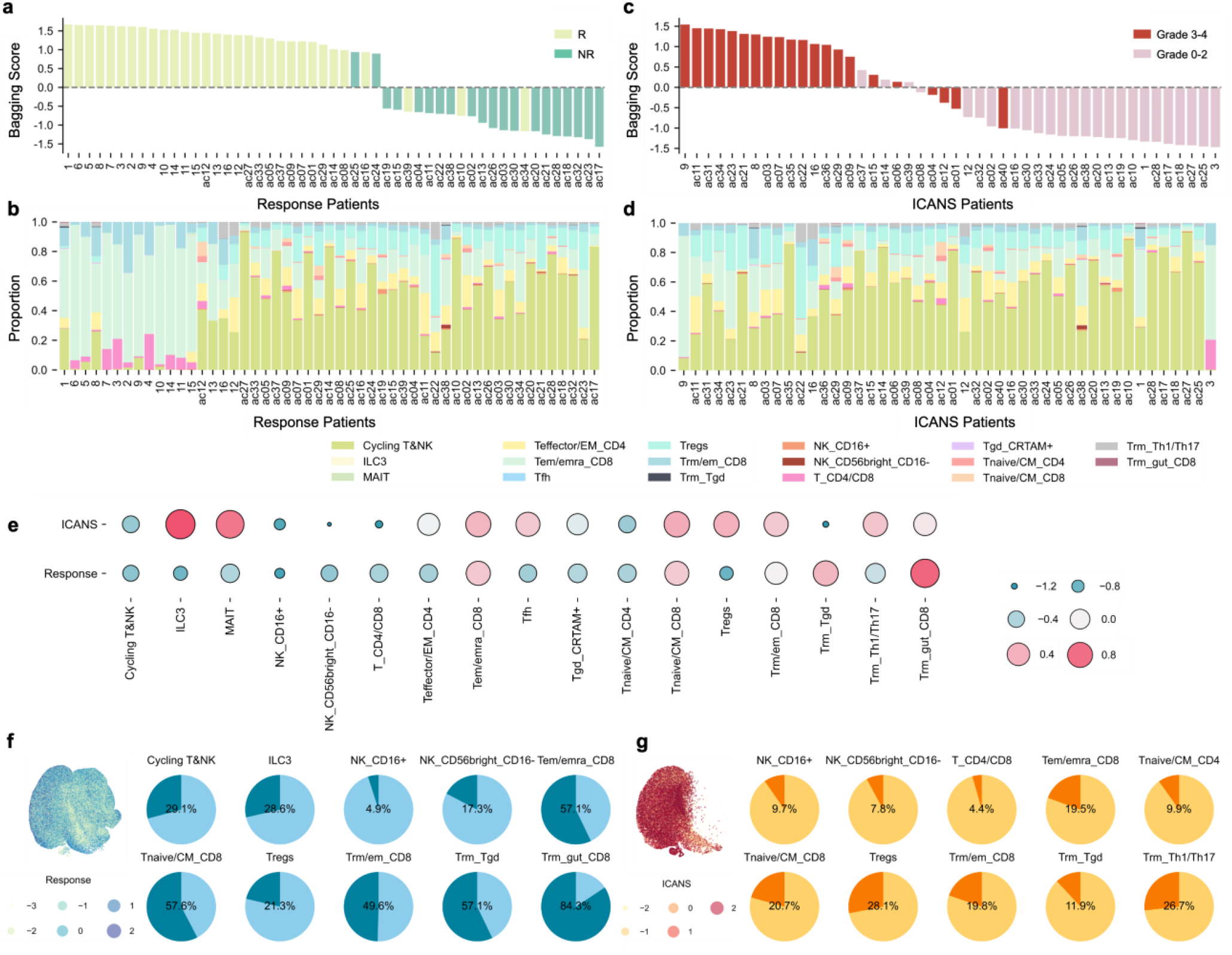
Evaluation of CART-GPT-response and CART-GPT-ICANS predictions on dual-labeled patients with cell-type-resolved model interpretation. a, Patient-level CART-GPT-response predictions. Each bar represents an individual patient, with the X-axis showing patient IDs and the Y-axis representing aggregated bagging scores. Light yellow-green bars indicate responders; teal bars indicate non-responders. b, T cell subtype composition across responders. Stacked bar plots show the TcellGPT-predicted cell type distribution for each response-labeled patient (X-axis = patients; Y-axis = proportion of each subtype). c, Patient-level CART-GPT-ICANS predictions. Aggregated bagging scores for ICANS severity prediction. The X-axis shows patient IDs; the Y-axis shows the bagging score. Dark red bars represent high-risk patients (Grade 3–4); lavender bars represent low-risk patients (Grade 0–2). d, T cell subtype composition across ICANS-labeled patients. Stacked bar plots display the cell type distribution across patients stratified by ICANS severity. e, Divergent model contributions across response and ICANS tasks. Bubble chart showing cell-type-specific contribution scores to ICANS prediction (top row) and response prediction (bottom row). Bubble size reflects total contribution; color encodes directionality (contribution to model). f, Contribution density in response-predictive cell types. Left: UMAP projection of cells colored by contribution to the response classification layer. Right: Pie charts show the proportion of cells within each TcellGPT-predicted subtype with contribution scores > 0. g, Contribution density in ICANS-predictive cell types. Left: UMAP projection of cells colored by contribution to the ICANS classification layer. Right: Pie charts show the proportion of cells within each TcellGPT-predicted subtype with contribution scores > 0.

Importantly, model interpretation revealed divergent sets of predictive features between the two tasks. For example, T_CD4/CD8 cells were more strongly associated with response prediction, whereas Teffector/EM_CD4 cells showed greater relevance for ICANS prediction (**Figure 4e**,**Supplementary Table 5**, p<0.05, Fisher’s exact test). Conversely, while Cycling T&NK cells were the most abundant population across nearly all patients, they did not substantially contribute to either model’s predictions. Rather than relying on bulk cell type abundance, CART-GPT selectively identifies functionally relevant subsets within each type. For instance, although cell types such as Tem/emra_CD8 and Tnaive/CM_CD8 were strongly associated with both treatment response and ICANS, only about 50% of these cells exhibited high contribution scores in the response task, and roughly 20% did so in the ICANS task (**Figure 4f-g, Extended Data Figure 5**). These results highlight CART-GPT’s ability to move beyond static cell type annotations and instead evaluate the functional relevance of individual cells in context, enabling more accurate, interpretable, and biologically meaningful patient-level predictions.

### CART-GPT identifies distinct gene programs underlying therapeutic response and ICANS severity

To further elucidate the immunological underpinnings of CART-GPT predictions, we investigated the model’s internal representation (embedding) layers to infer gene programs associated with therapeutic response. By analyzing the co-expression of genes involved in the embedding layers and inferring Gene Regulatory Network (GRN)^27^, we identified “metagenes”—clusters of co-regulated genes that collectively contribute to outcome prediction. These metagenes were grouped based on their inferred scores, allowing the extraction of functional gene modules implicated in treatment response or resistance (**Methods**). Visualization of these contribution-weighted metagenes suggested notable differences in transcriptional signatures between the patient groups with response (R) and no-response (NR) (**Figure 5a, Supplementary Table 6**). Interestingly, the 12_SCORE metagene appeared prominently enriched in responders and may reflect a cytotoxic and pro-inflammatory effector program. This module includes genes such as GZMA, GZMK, GNLY, CCL5, and KLRD1—associated with granule-mediated cytotoxicity, immune cell recruitment, and NK-like activity^28–30^—features that are generally linked to effective anti-tumor immune responses. Cell-type–specific expression analysis suggested that Tem/emra_CD8, Trm_Tgd, and Tem/em_CD8 subsets in responders may contribute to this cytotoxic transcriptional program, as many genes within the 12_SCORE metagene appeared to be expressed at higher levels in these populations (**Extended Data Figure 6**).

**Figure 5.**
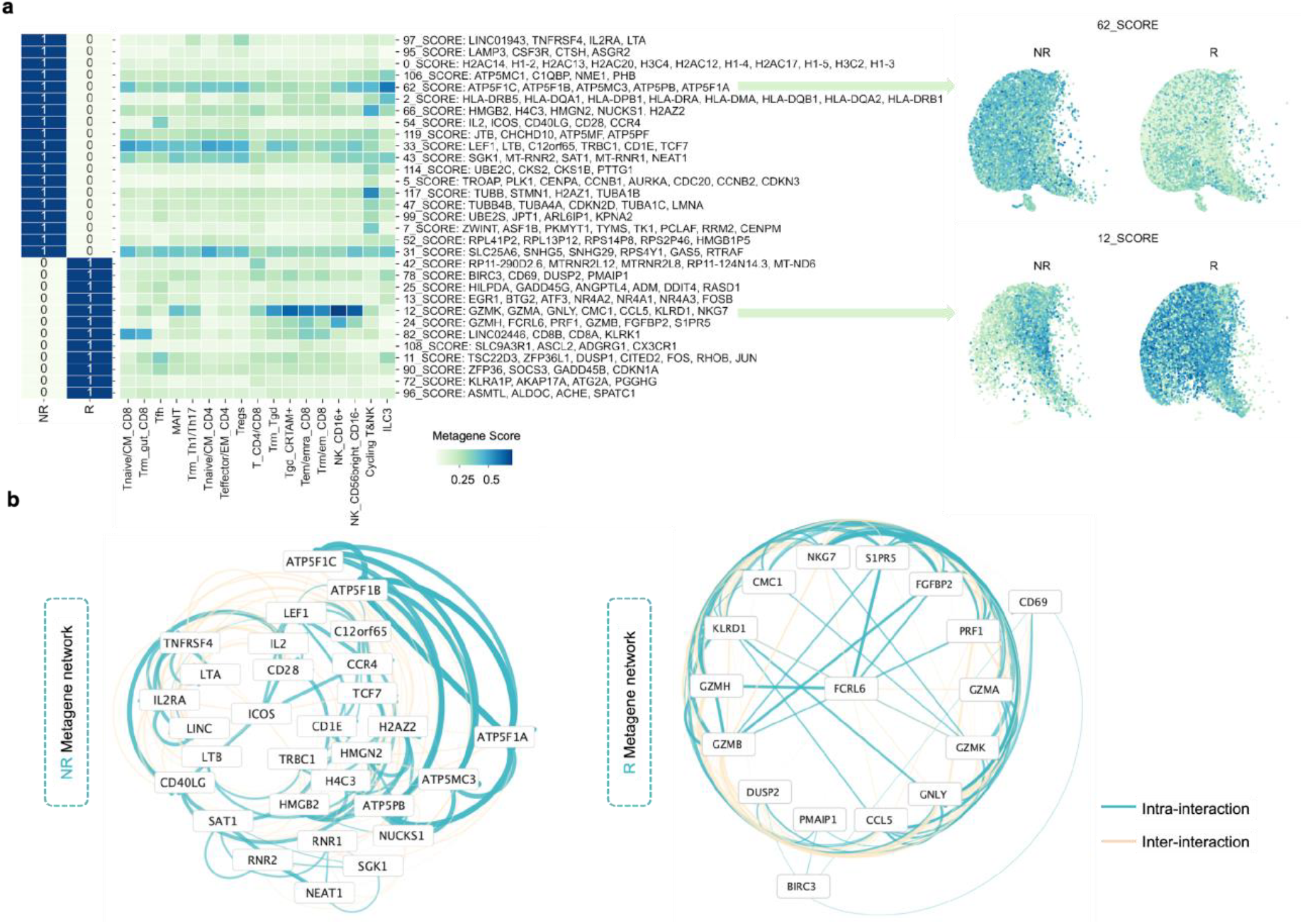
Therapy response–associated gene programs and regulatory networks revealed by CART-GPT. a, Metagene activity across responders and non-responders. Left: Heatmap showing the average CART-GPT metagene scores across annotated T cell subsets, stratified by response status. Each row corresponds to a metagene program; each column represents a T cell subtype. Right: UMAP projections of single cells colored by model-derived contribution scores for representative metagenes (12_SCORE and 62_SCORE). b, Regulatory networks in non-responders and responders. Left: Gene-gene interaction network inferred for non-responders; Right: corresponding network for responders. Nodes represent high-contribution genes, and edges indicate intra- and inter-metagene interactions, with edge thickness corresponding to interaction strength.

In contrast, the 31_SCORE, 33_SCORE, and 62_SCORE metagenes were more apparently enriched in non-responders. The 31_SCORE module includes transcripts associated with stress-adaptive or exhausted states (e.g., GAS5, RPS4Y1, SNHG5)^31,32^, while 33_SCORE includes genes linked to stem-like or memory-prone phenotypes (e.g., LEF1, TCF7, TRBC1)^33^. The 62_SCORE metagene, featuring mitochondrial respiratory genes (e.g., ATP5F1A, ATP5F1B, ATP5MC3)^34^, may point to altered metabolic activity. Notably, these gene programs appeared to be more selectively activated across specific cell types in non-responders (**Extended Data Figure 7**).

To further investigate the gene regulatory framework, we visualized metagenes and their GRNs using the network visualization software, Cytoscape^35^ (**Figure 5b, Extended Data Figure 8**).

The network of responders appeared more compact and modular, potentially reflecting a more focused transcriptional program. FCRL6 (Fc Receptor–Like 6)^36^, a molecule typically expressed by cytotoxic lymphocytes, emerged as a central hub, possibly indicating a coordinated regulatory role in cytolytic and pro-inflammatory signaling. In contrast, the network in non-responders was more diffuse and complex, with ICOS (Inducible T Cell Co-Stimulator, CD278) emerging as a hub gene and the most prominent interactions observed among mitochondrial genes such as ATP5F1A-C, ATP5PB, and ATP5MC3. Although ICOS is known to modulate T cell activation and differentiation^37^, its centrality—alongside the mitochondrial and memory-related gene modules—may reflect a functionally heterogeneous or metabolically altered immune state in non-responders.

Similarly, we applied the CART-GPT framework to explore gene programs potentially associated with ICANS severity. Inferred GRNs identified several metagenes that appeared more active in patients with high-grade ICANS (grade 3–4), including 31_SCORE, 65_SCORE, and 45_SCORE (**Figure 6a, Supplementary Table 7)**. These gene modules may reflect heightened inflammatory and immune activation states. For example, 65_SCORE includes genes such as CCL5 and IGHV7-81, which are linked to immune cell recruitment and B cell signaling^28^, while 45_SCORE features CD69 and NUCKS1, associated with T cell activation and chromatin remodeling^38,39^. Single-gene expression analysis showed that these metagenes were broadly expressed across various T cell subsets in the high-grade group (**Extended Data Figure 9**), potentially indicating a widespread and excessive immune response. In contrast, the 35_SCORE metagene was relatively enriched in low-grade ICANS (grade 0–2). This module includes genes such as SRSF2, SELL, and SPINT2, which are involved in mRNA splicing, leukocyte adhesion, and epithelial maintenance^40–42^. These genes appeared to be more selectively expressed across cell types (**Extended Data Figure 10**), which may suggest a more regulated and less inflammatory transcriptional landscape in patients with low ICANS.

**Figure 6.**
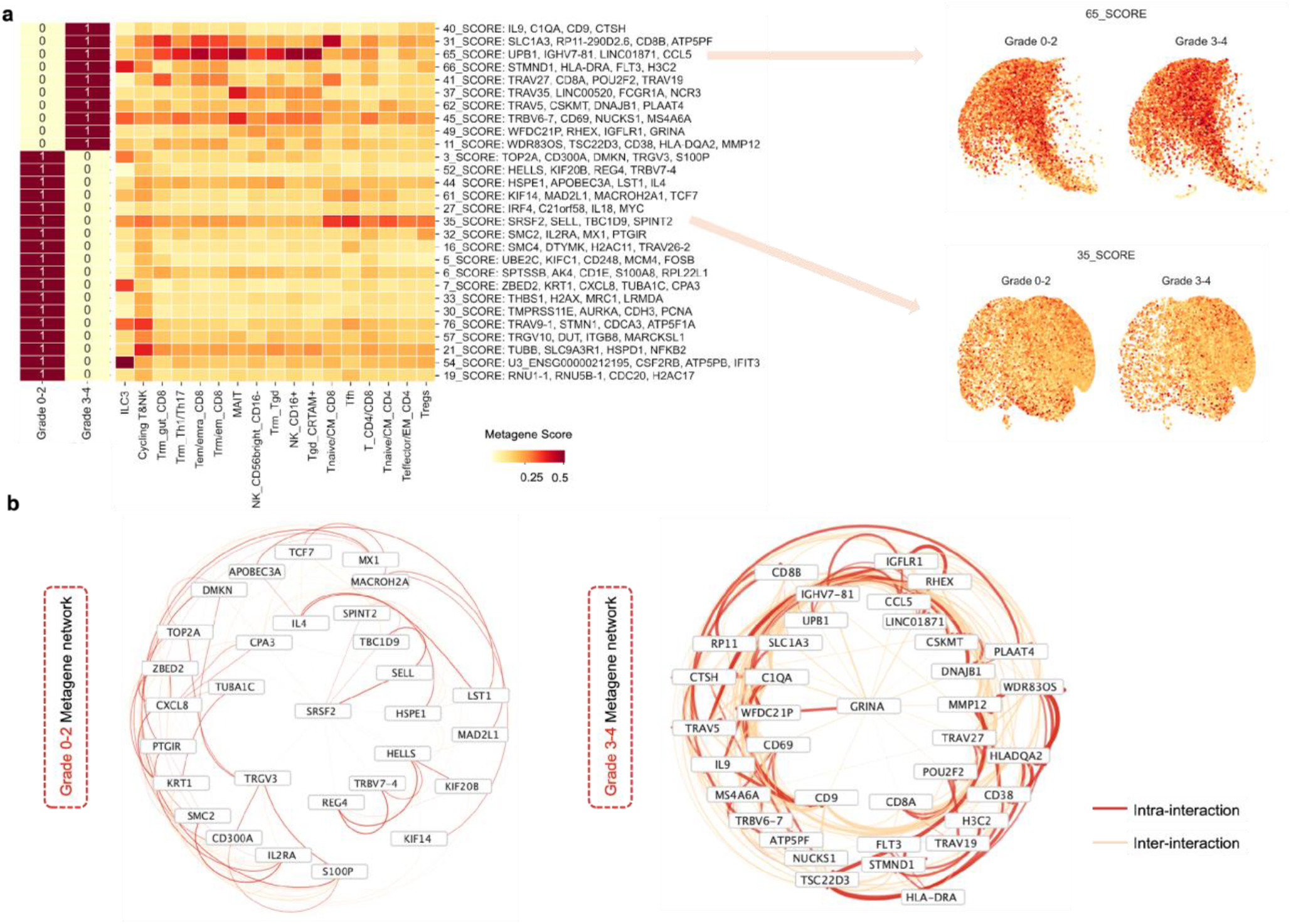
ICANS severity–associated gene programs and regulatory networks revealed by CART-GPT. a, Metagene activity across low-grade and high-grade ICANS groups. Left: Heatmap showing the average CART-GPT metagene scores across annotated T cell subsets, stratified by ICANS severity (Grade 0–2 vs. Grade 3–4). Each row corresponds to a metagene; each column represents a T cell subtype. Right: UMAP projections of single cells colored by model-derived contribution scores for representative metagenes (65_SCORE and 35_SCORE). b, Regulatory networks in low-grade and high-grade ICANS. Left: Gene-gene interaction network inferred for low-grade (Grade 0–2) ICANS patients; Right: corresponding network for high-grade (Grade 3–4) patients. Nodes represent high-contribution genes, and edges indicate intra- and inter-metagene interactions, with edge thickness corresponding to interaction strength.

Furthermore, we visualized the metagenes and GRNs to elucidate the potential distinct T cell languages that underlie low- and high-ICANS outcomes. High-grade ICANS may be characterized by a densely connected transcriptional architecture, with GRINA emerging as a central hub gene (**Figure 6b, Extended Data Figure 8**). GRINA, known for its role in calcium signaling and cellular stress resistance, has been implicated in neuronal survival and may represent a compensatory response to inflammation-associated neurotoxicity^43^. In contrast, the low-grade ICANS network displayed fewer and weaker gene-gene interactions, with SRSF2 identified as a possible hub. As a splicing regulator, the prominence of SRSF2 may indicate a more transcriptionally restrained or homeostatic profile^40^. These findings suggest that high-grade ICANS may be characterized by a more coordinated and pro-inflammatory transcriptional program, whereas low-grade ICANS may reflect a more regulated cellular state.

Notably, some gene-gene connections inferred by the model align with established immunological pathways, whereas others may reveal novel associations that warrant further investigation. Collectively, these findings suggest that CART-GPT encodes treatment outcomes through a latent T cell language, leveraging whole-transcriptomic gene programs, some of which may remain unexplored. Thus, AI-based modeling may also serve as a valuable tool for generating mechanistic hypotheses about CAR-T cell function and therapeutic outcomes.

## Discussion

We developed CART-GPT, the first foundation model-driven framework fine-tuned on single-cell transcriptomic data specifically for predicting patient-specific therapeutic outcomes in CAR-T therapy. Although CAR-T therapy has become a cornerstone of treatment for certain hematologic malignancies^1,4^, its clinical implementation continues to face critical bottlenecks—particularly in stratifying patients by therapeutic response or toxicity risk^7,10^. CART-GPT addressed this critical gap by decoding the immune signatures associated with efficacy or neurotoxicity into a T cell language.

Importantly, because most CAR-T cells are derived autologously, CART-GPT is uniquely positioned to infer both the intrinsic immune status of the patient and the functional state of the infused CAR-T cells. This dual-layer perspective captures how host immunity and CAR-T cell fitness together shape therapeutic outcomes. Its modular design, transformer-based modelling, transfer learning approach, and innovative cell bagging strategy enable robust patient-level predictions from noisy, high-dimensional scRNA-seq data. Moreover, CART-GPT offers underlying insights into the collective behavior of T cells, interpreting outcomes as an emergent property of immune system dynamics. On held-out independent patient data, the model achieves an AUC of ~0.8, indicating strong, clinically meaningful performance and promising potential for future commercial applications. These results not only enhance our understanding of CAR-T biology but also set the stage for precision-guided interventions and more informed clinical decision-making in CAR-T therapy.

Although foundation models hold great promise for single-cell analysis by providing general-purpose representations of cellular states^17–19^, their direct utility in clinical applications remains limited without careful task-specific fine-tuning. In the context of CAR-T therapy, clinical prediction tasks—such as response and toxicity stratification—require more than general transcriptional embeddings; they demand highly specialized adaptation to disease- and treatment-specific immunological contexts. Such fine-tuning is only feasible with access to high-quality, well-annotated datasets. A central contribution of this study is the construction of a comprehensive and curated CAR-T cell atlas comprising over 1.12 million single cells from patients with detailed clinical outcomes, including treatment response and ICANS severity.

Importantly, we applied rigorous batch correction to integrate heterogeneous datasets into a biologically coherent, uniform atlas suitable for downstream modeling. This dataset provides the critical foundation for effective model specialization, enabling CART-GPT to learn subtle, clinically relevant transcriptional patterns. The scale, resolution, and outcome-linked nature of the atlas make it uniquely suited for training robust, generalizable models that bridge cellular heterogeneity with patient-level prediction, highlighting the indispensable role of high-quality data in the success of AI-driven precision medicine.

While CART-GPT demonstrates strong predictive capabilities, several limitations persist. The current model is trained primarily on CD19 CAR-T datasets, which may constrain its generalizability to other CAR targets. Expanding CART-GPT to incorporate data from diverse CAR constructs and disease contexts will be critical for broader clinical applicability.

Additionally, as more datasets that will become available, there is potential to enhance the granularity of ICANS prediction—moving beyond binary classification (grade 0–2 vs. grade 3–4) toward a more refined stratification across the full ICANS grading spectrum (grades 0 to 4).

Moreover, while CART-GPT offers a practical tool for outcome prediction, it encodes complex, context-specific immune dynamics that go beyond current biological understanding. Although we employed model interpretation techniques to uncover latent gene programs and regulatory networks, much of the model’s decision-making remains a black box. Thus, CART-GPT not only serves as a predictive engine but also as a hypothesis-generating framework, offering a starting point for deeper mechanistic investigation.

Ultimately, CART-GPT illustrates the promise of large language models in decoding immune cell behavior, although unlocking their full potential will depend on continued integration of computational predictions with experimental validation and clinical interpretation. Beyond CAR-T therapy, the CART-GPT framework is inherently scalable and generalizable to other immunotherapies and diseases characterized by complex cellular ecosystems. Its modular architecture and foundation model backbone allow for efficient adaptation to new clinical contexts, provided appropriately curated single-cell atlas is available. The combination of transfer learning, cell state modeling, and outcome-level aggregation makes the framework broadly applicable to tasks such as predicting response to immune checkpoint inhibitors or characterizing immune dysregulation in autoimmune or infectious diseases. As such, CART-GPT provides a blueprint for extending large-scale single-cell modeling to diverse areas of precision immunology.

## Methods

### Data Collection

To construct a foundation for modeling T cell phenotypes and predicting clinical outcomes, we first assembled a large reference T cell atlas derived from publicly available single-cell RNA-sequencing (scRNA-seq) datasets hosted on the CELLxGENE database^20^. This reference atlas spans 17 human tissues—including immune-relevant compartments such as peripheral blood, bone marrow, kidney, spleen, and lymph nodes—and captures a broad spectrum of T cell subtypes, including CD4^+^ helper T cells, CD8^+^ alpha-beta memory T cells, gamma-delta T cells, regulatory T cells, and other functionally distinct subsets, providing a comprehensive landscape of natural T cell biology.

In addition, we manually curated a large CAR-T cell dataset from published studies, compiling data from both main figures and supplementary materials. All CAR-T cells were sampled from infusion products (IPs), specifically the residual cells collected by washing standard-of-care axi-cel infusion bags after administration. These IP-derived cells are particularly valuable because they retain transcriptomic signatures reflective of the patient’s post-treatment clinical trajectory. The dataset is annotated with rich clinical metadata, including binary and graded labels for treatment response and immune effector cell–associated neurotoxicity syndrome (ICANS, graded 0–4).

For quality control, we applied standard filtering steps: (1) genes expressed in fewer than three cells were removed, and (2) cells expressing fewer than 100 genes were excluded. To correct for batch effects arising from technical or donor variability, we applied scVI (single-cell variational inference)^44^, a deep generative model that learns latent representations of gene expression while integrating across datasets. The curated CAR-T dataset includes 782,558 cells from 199 patients annotated with treatment response labels, and an additional 343,061 cells from 56 patients annotated with immune effector cell–associated neurotoxicity syndrome (ICANS) outcomes (**Supplementary Table 1**). All datasets were processed into unified AnnData objects using Scanpy, with layers for raw counts, normalized expression, and scVI-corrected embeddings^44^. Metadata fields include T cell subtype annotations, patient identifiers, treatment response status, and ICANS grades.

### CART-GPT

CART-GPT is an all-in-one AI toolbox designed for T cell subtype annotation and for predicting CAR-T therapy outcomes, including treatment response and ICANS risk. It leverages large language models (LLMs) to decode the cellular landscape of infusion product (IP) CAR-T cells and uncover potential links to clinical efficacy and toxicity. For model development, we applied transfer learning to adapt and fine-tune a foundation LLM model, scGPT, and further introduced a novel cell bagging strategy to identify high-impact cellular subpopulations that drive patient-level predictions.

#### LLM architectures for CART-GPT models

CART-GPT models were re-trained from the scGPT model through transfer learning^19^. We freeze the gene embedding parameters and sequentially fine-tune the remaining model parameters to build three task-specific models: 1) TcellGPT is fine-tuned from scGPT for general T cell annotation. 2) CART-GPT-response is fine-tuned from TcellGPT to predict patient therapy response. 3) CART-GPT-ICANS are subsequently fine-tuned from CART-GPT-response for ICANS prediction. The parameters for embedding layers, multiple head in the transformer layers, layer neurons, and predictive labels for the classification layers can be found in **Supplementary Table 8**.

The architecture of the model is described below

a. Input Representation (Gene Embeddings): Let *x*_*i*_ represent the expression level of the *i*-th gene in a single cell. The gene expression vector for a cell is:

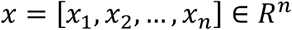

where *n* is the number of genes. Each gene expression value *x*_*i*_ is converted into a gene embedding vector using an embedding function *E*(*x*_*i*_) ∈ *R*^*d*^, where *d* is the embedding dimension. The resulting embedded representation of the gene expression profile is:

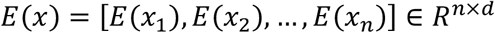
b. Transformer Architecture: CART-GPT, like scGPT, utilizes transformer layers with multi-head self-attention and embedding and transformer dimensions to model interactions between genes. The self-attention mechanism computes attention scores for gene pairs, allowing the model to learn complex gene–gene dependencies. Specifically, the architecture and related parameters used in this study are shown in Figure 2a and Figure 3a. For a given input embedding *E*(*x*), the attention mechanism computes attention weights *A*_*ij*_ for each gene pair (*i, j*) as:

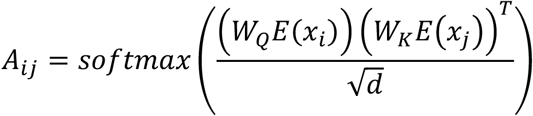

where *W*_*Q*_ and *W*_*K*_ are the query and key projection matrices, and *d* is the dimensionality of the embeddings. The attention-weighted sum of values *V*_*j*_ (from value projection *W*_*V*_) gives the updated embedding for each gene:

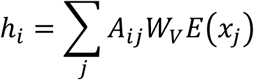

where *h*_*i*_is the output for gene *i*.
c. Positional Encoding: To account for the position of genes, a positional encoding P ∈ *R*^*n*×*d*^ is added to the embeddings:

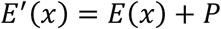

This helps the model understand gene order or structural relationships in the input data.
d. Prediction: The output of the Transformer layers is a new set of representations for the gene expression profile:

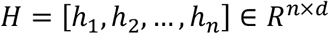

These representations are processed by multiple fully connected layers (n=17 for cell types and n=2 for CAR-T cell therapy outcomes and risks) and a softmax function for predictions:

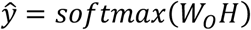

where *ŷ* is the predicted probability distribution.

### Transfer learning for CART-GPT models

We set the weights of the pre-trained model.

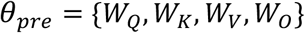

where: *W*_*Q*_, *W*_*K*_, *W*_*V*_ are the query, key, and value weight matrices in the self-attention mechanism, and *W*_*O*_ is the output weight matrix.

The loss function used during pre-training is denoted as *L*_*pre*_, and it depends on the pre-trained model’s weights *θ*_*pre*_.

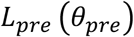

Let *θ*_*new*_ represent the weights of the model after fine-tuning on a new dataset *D*_*new*_.

The new loss function is:

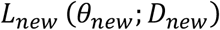

where *D*_*new*_ is the dataset being used for fine-tuning, and *θ*_*new*_ represents the weights adapted during fine-tuning.

The goal of fine-tuning is to minimize the new loss function *L*_*new*_ with respect to the new weights *θ*_*new*_:

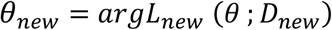

During training, the weights are updated using gradient descent:

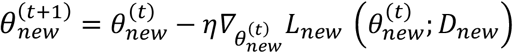

where:

- 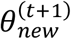 is the weight at the t-th iteration,
- η is the learning rate,
- 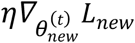 is the gradient of the loss function with respect to 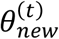.

The fine-tuning process involves minimizing the new loss function iteratively until convergence:

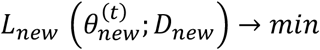

Each fine-tuning step on a new dataset enables the model to acquire new task-specific knowledge, which is encoded in the updated parameters *θ*_*new*_ in the GPT model by

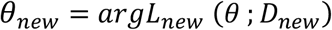

We denote the T cell scRNA-seq datasets by their associated metadata as *D*_*Tcells*_, *D*_*Response*_, *D*_*ICANS*_. For patient datasets, we randomly split each of them into 5-fold or 3-fold training and testing datasets at the patient level. For ICANS labeling, grades 0–2 were categorized as ICANS-low, while grades 3-4 were categorized as ICANS-high.

Based on the pre-trained model’s weights *θ*_*pre*_ from a fine-tuned model of scGPT for cell type annotation, we applied fine-tuning to each of these datasets.

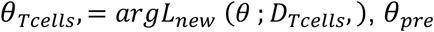

The model with *θ*_*Tcell*_ is defined as TcellGPT, where *L*_*new*_ is defined as the difference between predicted T cell subtypes and the annotated T cell subtypes in the *D*_*Tcells*_.

Based on the weights *θ*_*Tcells*_ from TcellGPT for T cell subtype annotation, we applied fine-tuning to *D*_*Response*_.

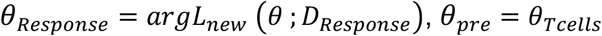

The model with *θ*_*Response*_ is defined as CART-GPT-response. Unlike models that predict cell type, CART-GPT-response predicts the patient’s therapy response status. During training, *L*_*new*_ is defined as the difference between the predicted cell label *ŷ* and the clinical information 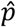 of response.

Finally, based on the weights *θ*_*Response*_ from CART-GPT-response for T patient therapy response, we applied fine-tuning to *D*_*ICANS*_ to build models for predicting ICANS, respectively.

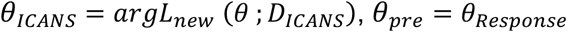

The model with *θ*_*ICANS*_ is defined as CART-GPT-ICANS. Here, *L*_*new*_ is defined as the difference between the predicted cell label *ŷ* and the clinical information 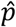 for ICANS status.

### Patient-Level Score Aggregation from Cell-Level Predictions

To infer patient-level clinical status (e.g., treatment response or ICANS) from cell-level predictions (i.e., classification layer outputs), we developed a strategy that groups cells based on classification layer outputs and aggregates their classification-layer output values into a patient-level predictive score. We refer to this approach as cell bagging, which enables outcome predictions at the patient level by leveraging single-cell information. Cell bagging involves three main steps: (i) excluding cells that cannot be classified by the CART-GPT output layers; (ii) identifying critical cell types in classification layers; (iii) bagging the cells from steps (i) and (ii).

In step (i), results from transfer learning revealed that certain cells show close predictive probabilities (*ŷ* = *softmax*(*W*_*O*_ *H*)). This might be a reality that not all cells in the IPs take part in determining the outcomes and risks of interest. To exclude the cells with weak associations with response or ICANS, CART-GPT-response and CART-GPT-ICANS used a criterion defined by the outputs in the classification output layer (n=2, for negative and positive, respectively),

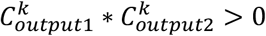

where *k* = 1, 2, ⋯, *K* is the cell ID. 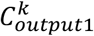 and 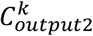 are the output values of cell *k* in the output layer.

In step (ii), results from transfer learning revealed that not every cell type takes part in determining the outcomes and risks to the same extent. To quantify the extents for cell types, we aggregated the softmax values (*ŷ*) of classification layer outputs. The aggregated softmax values 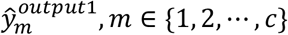 and 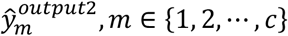 for cell types obtain the critical cell types for positive and negative status for response or ICANS.

In step (iii), for each patient (*p*), we define the predictive scores for response or ICANS using cell bagging. The score *S*_*p*_ is defined as below:

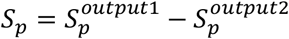

where *S*_*p*_ is the score to infer patient-level clinical status, 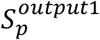 is the score inferred from the cell bagging for classification output 1 (positive), 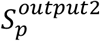 is the score inferred from the cell bagging for classification output 2 (negative),

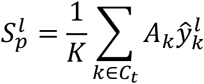

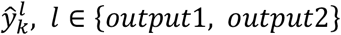 is the softmax probability of cell *k* predicted by classification output node *l*, the number of the cells participating bagging process is 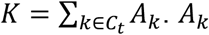 denotes the learning weight evaluating whether this cell has a strong association with the predictive statuses (as defined in step (i)), defined as below:

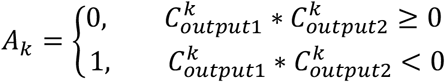

and *C*_*t*_ denote the ranked cell type clusters, selected based on the average cell-level score within each cluster (as defined in step (ii)). The ranking criterion is defined as follows:

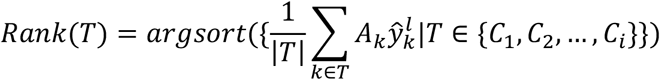

where *C*_1_, *C*_2_, *⋯, C*_*i*_ represent cell types derived from a single patient. Based on the learned weights, cell type clusters ranked higher by the above equation are interpreted as contributing more strongly to positive predictions (response or high ICANS risk), while lower-ranked clusters are associated with negative predictions (non-response or low ICANS risk). To reduce ambiguity and potential noise from mid-ranked clusters with uncertain contribution, we selected the top 3 ranked cell types (with cell number > 100) from each classification layer for patient-level score aggregation.

We converted a probability output from sigmoid values for each patient to a signed confidence score (e.g., confidence for positive vs negative class). Thus, each patient has either a positive value to predict positive status or a negative value to predict negative status. This is applicable to 2 CART-GPT models for response and ICANS.

### Gene Regulatory Network Inference and Visualization

To infer gene regulatory networks (GRNs) and associated gene programs^27^, we utilized the gene embedding-based GRN inference module. Gene embeddings were extracted from the output layer of the fine-tuned CART-GPT-response model or CART-GPT-ICANS model, and a nearest-neighbor graph was constructed to define gene–gene relationships based on embedding similarity. From this graph, metagenes representing co-expressed gene programs were identified. To visualize these regulatory networks, the inferred GRN data were imported into Cytoscape (v3.10.3)^35^. Networks were rendered using both circular and radial layouts, with nodes representing genes and edges denoting predicted regulatory interactions.

### Data analyses and model benchmarking

The pre-trained scGPT model (v0.2.1)^19^ was fine-tuned on our curated datasets to generate three task-specific models: TcellGPT, CART-GPT-response, and CART-GPT-ICANS. Model training was GPU-accelerated using CUDA (v11.7) and optimized with FlashAttention (v1.0.4) to enhance Transformer efficiency.

Model performance and patient-level predictions were evaluated using receiver operating characteristic (ROC) curves and the area under the ROC curve (AUC), implemented via scikit-learn (v1.2.2). The difference in outcome classification between CART-GPT tools and other approaches was calculated by Adjusted Rand Index (ARI)^45^. For benchmarking, UMAP projections were generated in Scanpy using the Leiden algorithm, and T cell subtype annotations were performed using CellTypist (v1.6.3) with the Immune_All_Low.pkl reference model19^21^.

All figures were generated using Python (v3.10.14) with matplotlib (v3.8.4), seaborn (v0.13.2), scikit-learn (v1.2.2), and Scanpy (v1.10.0). The predictive heatmaps were visualized using the scanpy.pl.matrixplot function. UMAP visualizations were rendered with built-in Scanpy plotting tools. AUC and ARI metrics were computed using sklearn.metrics.

## Data availability

Atlas-scale T-cell and CAR-T cell scRNA-seq data are publicly available at CELLxGENE and the GEO database^20^. The CAR-T cell atlas used in this work was uploaded to GitHub.

## Code availability

The CART-GPT package is available at GitHub https://github.com/guangxujin/CART-GPT.

## Author contribution

G.J. and Y.L. conceived and designed the study. W.G., Z.J., R.J., X.L., and Y.Z., collected the atlas data and clinical information. T.T.M., X.S., and G.J., conducted the data analysis. T.J., X.S., and G.J., contributed to modeling and software. T.T.M, X.S., T.J., T.M., and G.J., contributed to results and figures. G.J. supervised the project. T.T.M., X.S., T.J., Y.L., and G.J. wrote the manuscript with input from all authors. All authors reviewed and approved the final version of the manuscript.

## Acknowledgements

This work was supported by a start-up fund from Atrium Health Wake Forest Baptist (G.J). We also acknowledge support of the Wake Forest Baptist Comprehensive Cancer Center Bioinformatics Shared Resource, supported by [P30CA012197]. The content is solely the responsibility of the authors and does not necessarily represent the official views of the National Cancer Institute. This work is supported in part by the Digital Health and Geospatial Analytics Program (X.S) from National Research Council Canada. This research was enabled by the computational resources provided in part by the Digital Research Alliance of Canada (the Alliance) (https://alliancecan.ca/).

## Extended Data Figures

**Extended Data Figure 1.**
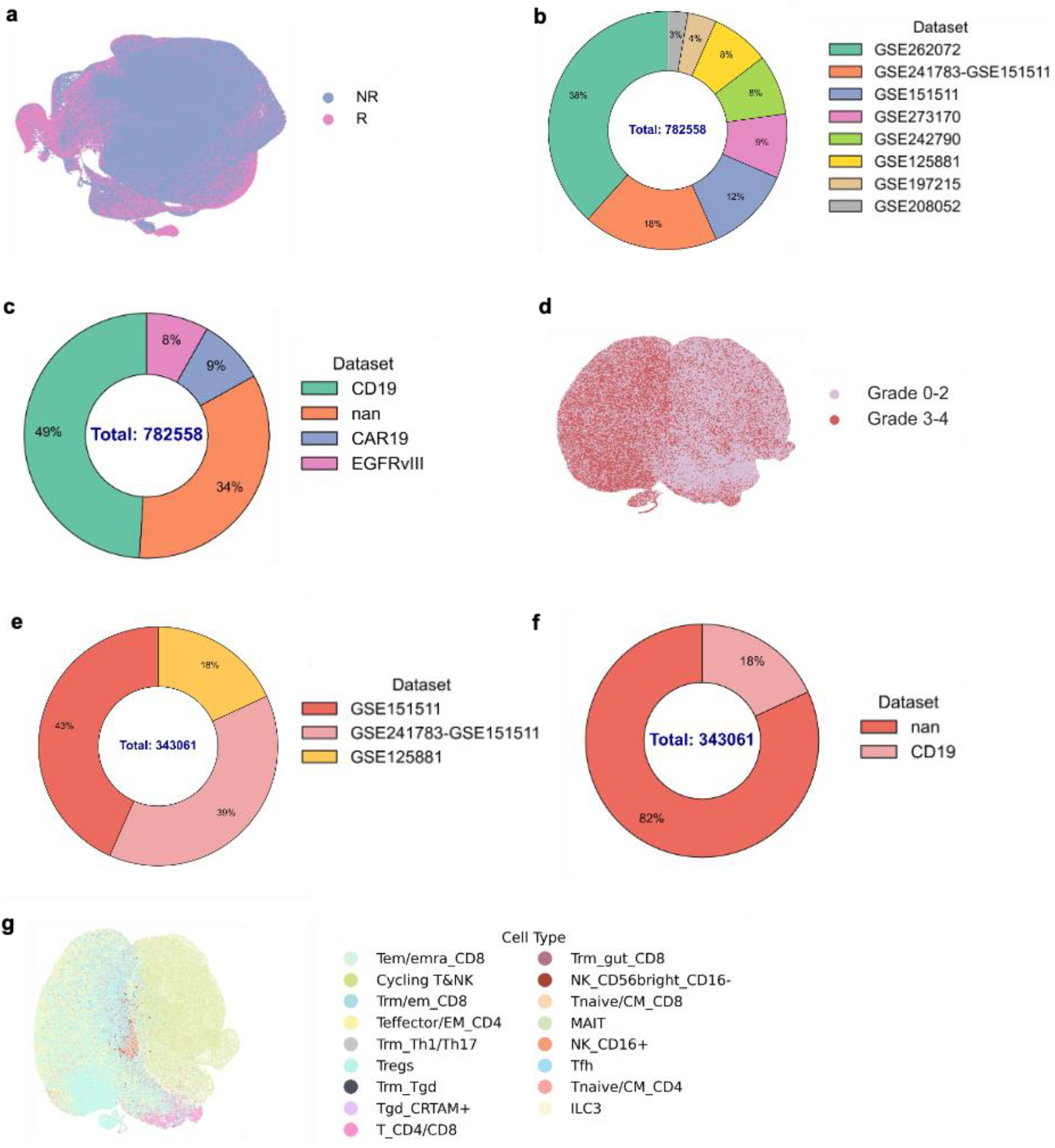
Overview of the CAR-T cell atlas. a, d, g, UMAP projections of all CAR-T cells stratified by clinical outcomes. a, Cells colored by treatment response (NR: non-responder, R: responder). d, Cells colored by ICANS severity (Grade 0–2 vs. Grade 3–4). g, T cell subtype predictions for all CAR-T cells using TcellGPT. b, e, Donut charts summarizing dataset composition. b, response datasets. e, ICANS datasets. c, f, Donut charts showing CAR-T construct types within response (c) and ICANS (f) datasets.

**Extended Data Figure 2.**
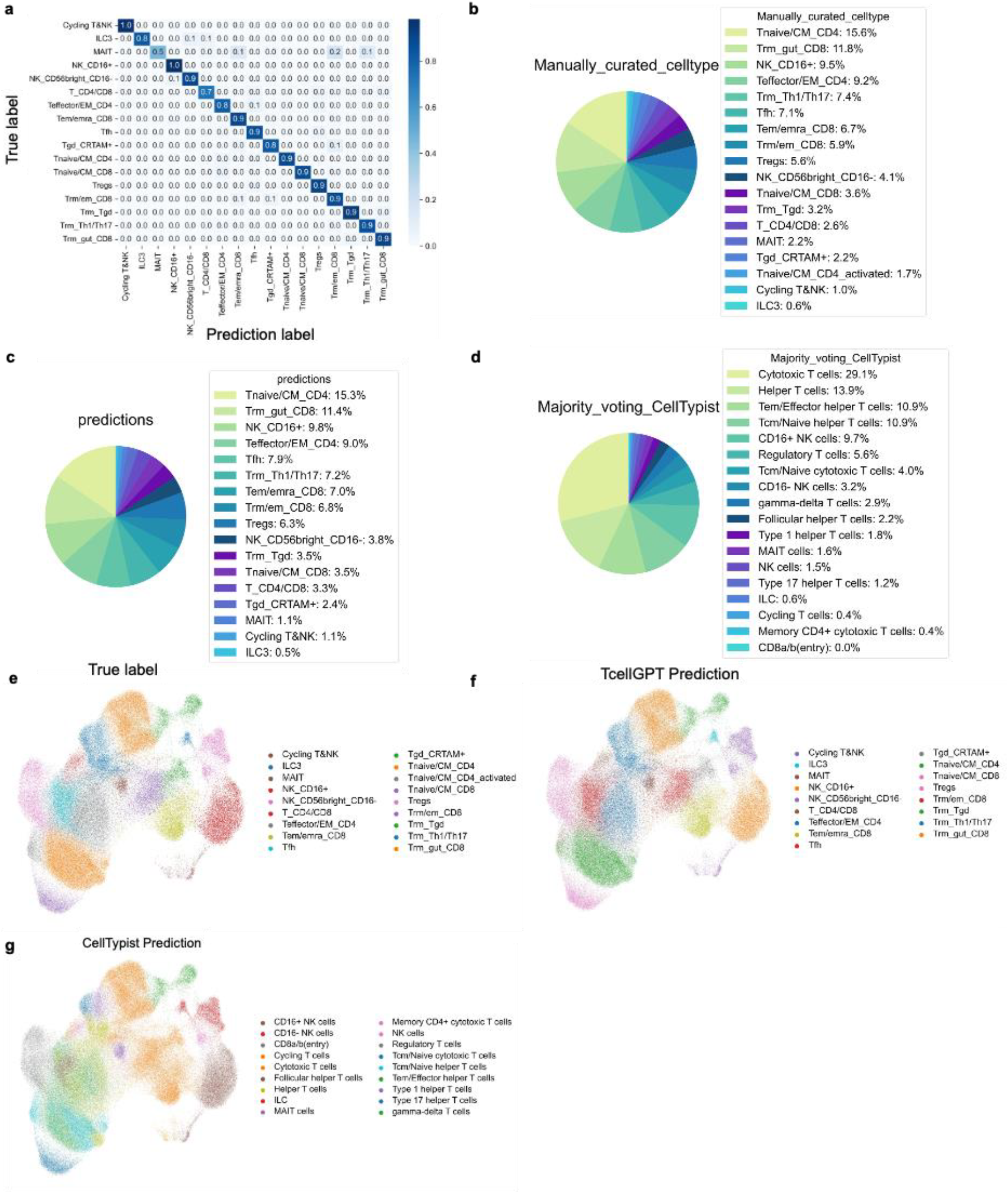
Evaluation of TcellGPT performance in T cell subtype annotation. a, Heatmap comparing TcellGPT-predicted T cell subtypes with manually curated labels across 17 cell types. b–d, Pie charts showing the distribution of T cell subtypes based on: b, manual annotations; c, TcellGPT predictions; and d, CellTypist majority-voting predictions. Legends are ordered by relative cell type abundance. e–g, UMAP projections of cells colored by: e, manual annotations; f, TcellGPT predictions; and g, CellTypist predictions.

**Extended Data Figure 3.**
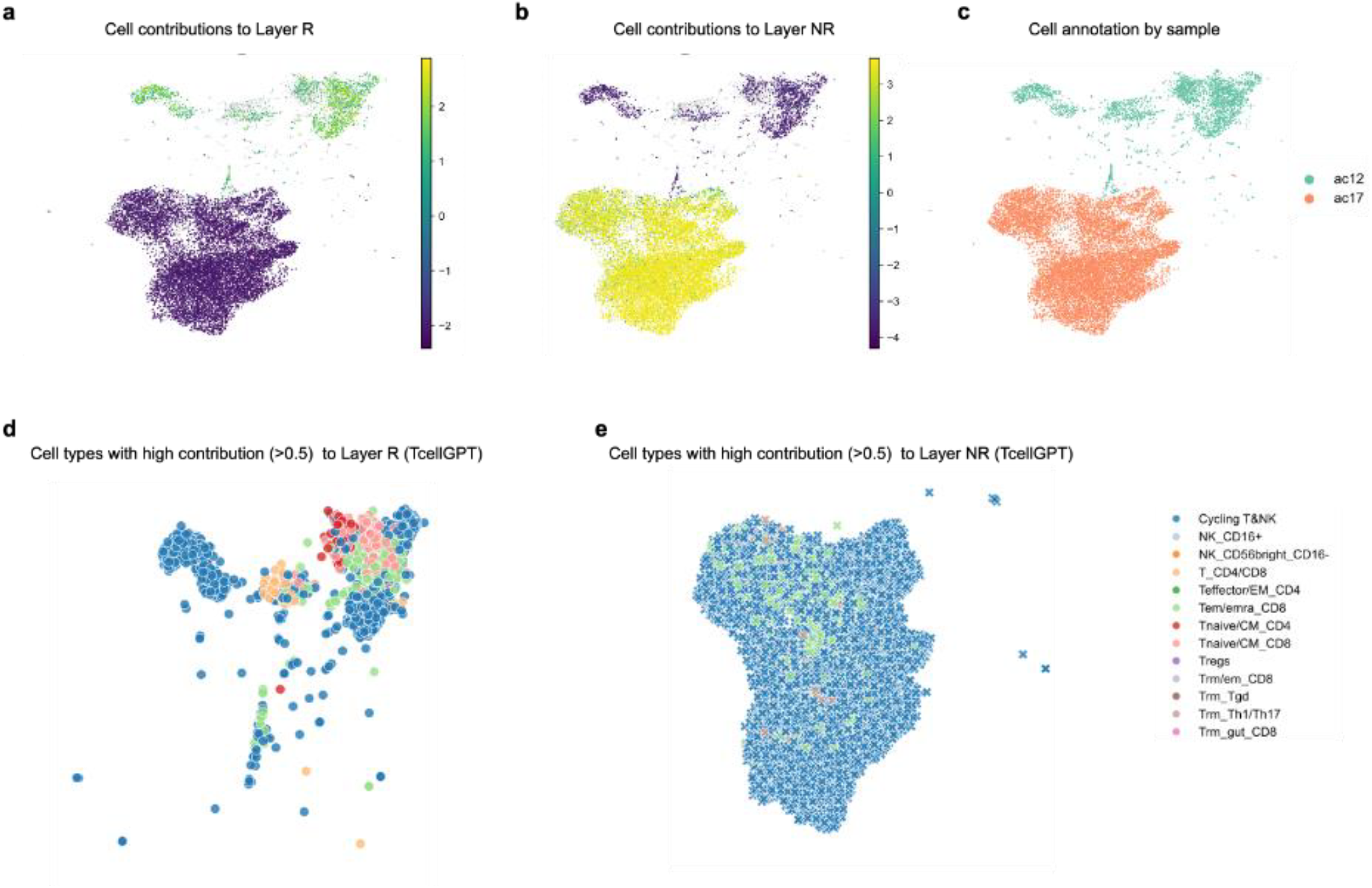
Patient-level visualization of CART-GPT-response annotations for representative patients. a–b, UMAP projections of cells from two representative patients, ac12 (responder) and ac17 (non-responder), colored by their contribution scores to the responder (R) and non-responder (NR) classification layers, respectively. c, UMAP projection of the same cells colored by patient sample, highlighting ac12 and ac17. d–e, Visualization of the top contributing cell types (contribution value > 0.5) driving classification outcomes in each patient. Cells are colored according to their TcellGPT-predicted identities.

**Extended Data Figure 4.**
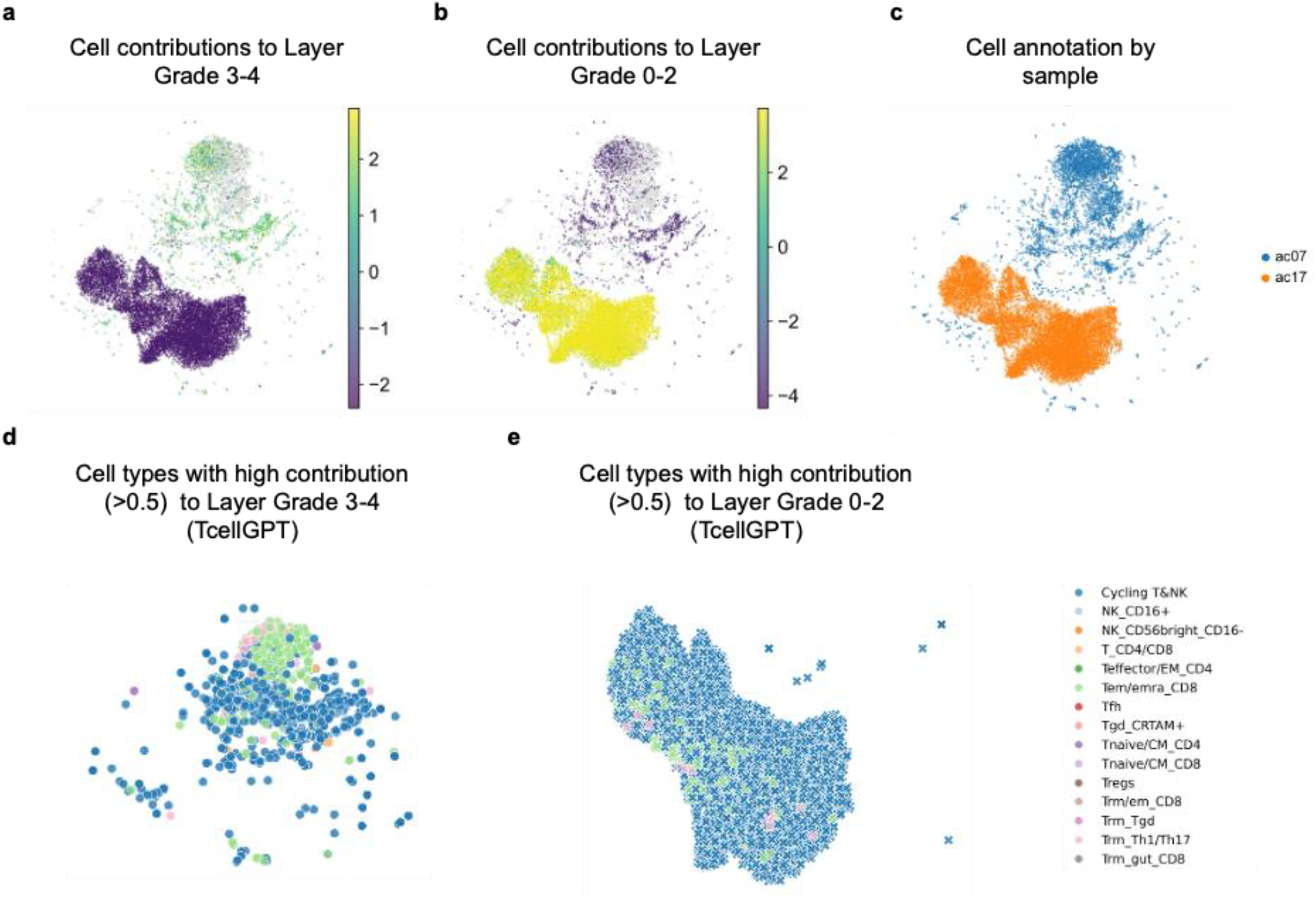
Patient-level visualization of CART-GPT-ICANS annotations for representative patients. a–b, UMAP projections of cells from two representative patients, ac07 (Grade 3–4) and ac17 (Grade 0– 2), colored by their contribution scores to the Low and High ICANS classification layers, respectively. c, UMAP projection of the same cells, colored by patient sample, highlighting ac07 and ac17. d–e, Visualization of the top contributing cell types (contribution value > 0.5) driving classification outcomes in each patient. Cells are colored according to their TcellGPT-predicted identities.

**Extended Data Figure 5.**
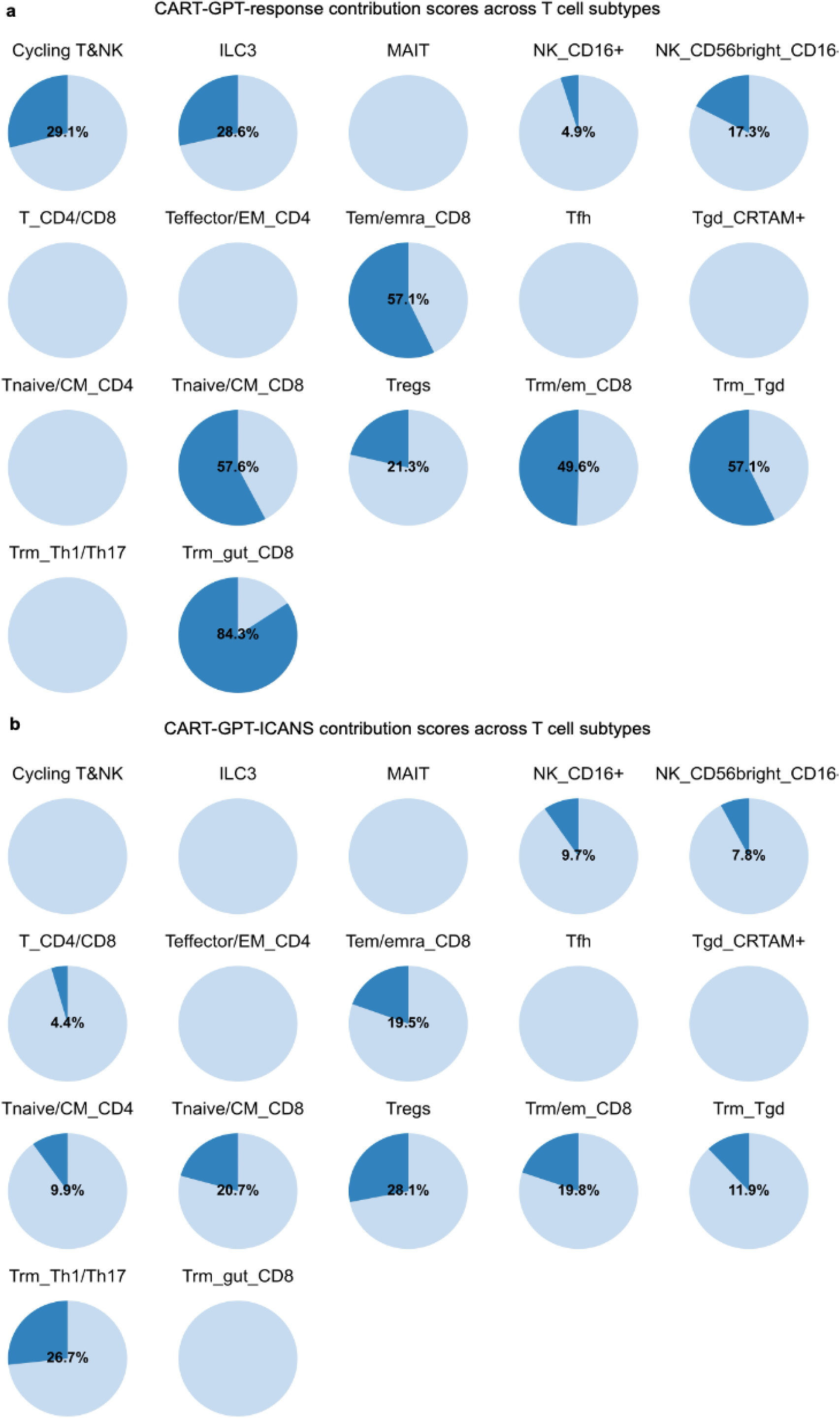
Proportion of cells with high CART-GPT contribution scores across T cell subtypes. a. Pie charts show the percentage of cells within each TcellGPT-predicted subtype that exhibit a contribution score greater than 0 to the CART-GPT-response model. Dark blue segments represent the fraction of high contributing cells, while light blue segments indicate low contributing cells. b. Pie charts show the percentage of cells within each TcellGPT-predicted subtype that exhibit a contribution score greater than 0 to the CART-GPT-ICANS model. Dark blue segments represent the fraction of high contributing cells, while light blue segments indicate low contributing cells.

**Extended Data Figure 6.**
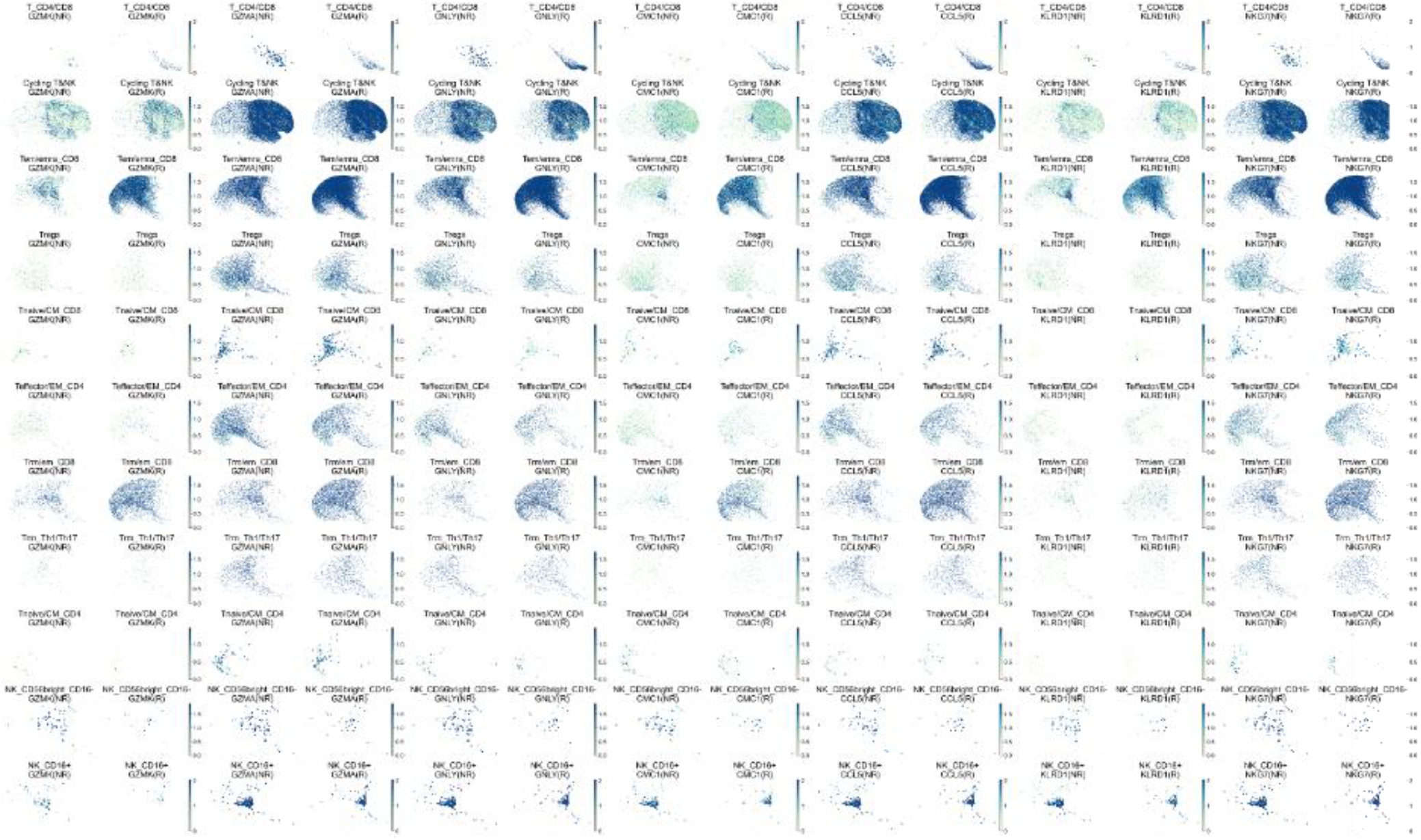
Cell-type–resolved expression of representative genes from the 12_SCORE metagene. Color intensity indicates gene expression, with darker shades reflecting higher expression. Each row represents a distinct cell subtype, and each column corresponds to a gene.

**Extended Data Figure 7.**
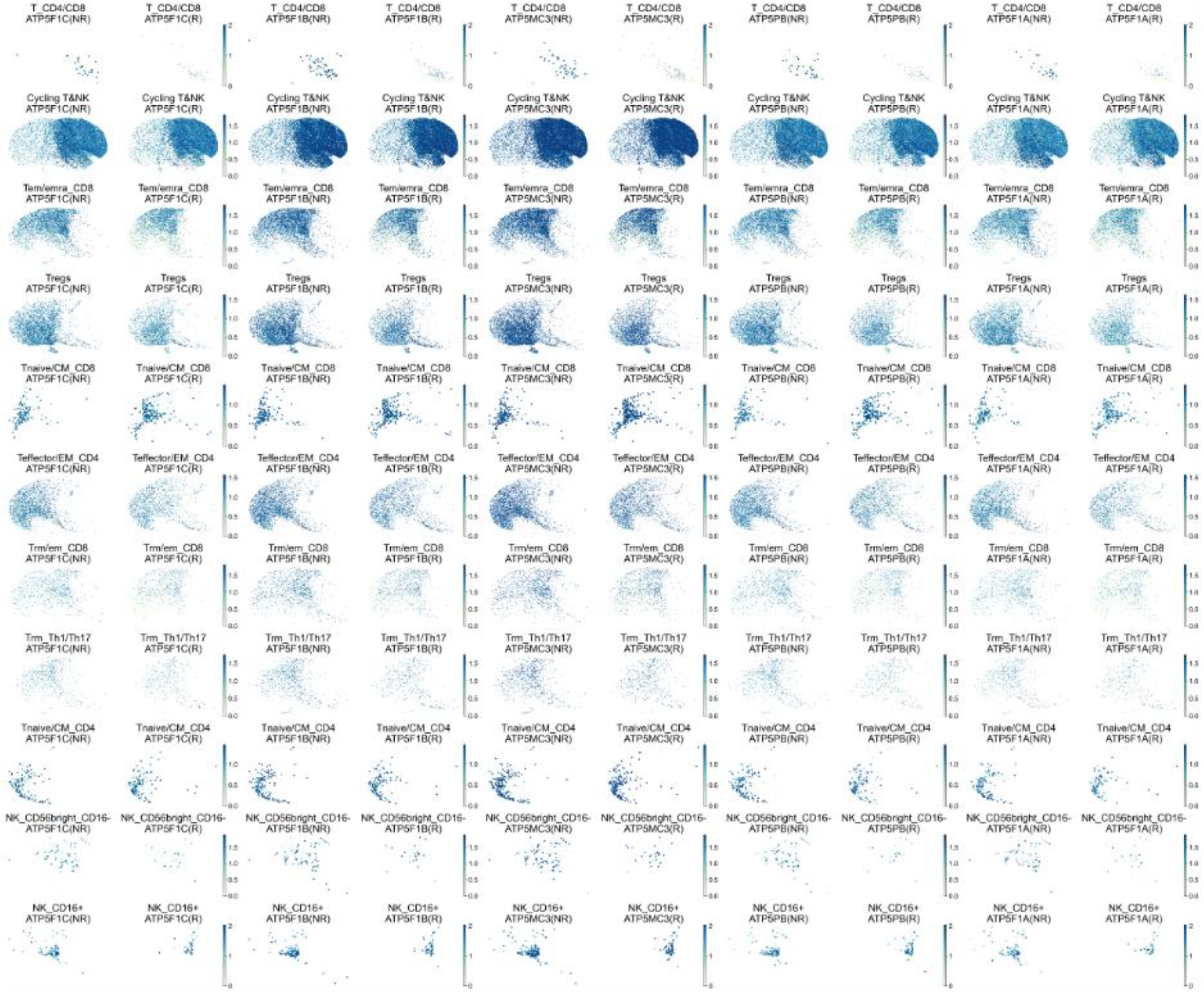
Cell-type–resolved expression of representative genes from the 62_SCORE metagene. Color intensity indicates gene expression, with darker shades reflecting higher expression. Each row represents a distinct cell subtype, and each column corresponds to a gene.

**Extended Data Figure 8.**
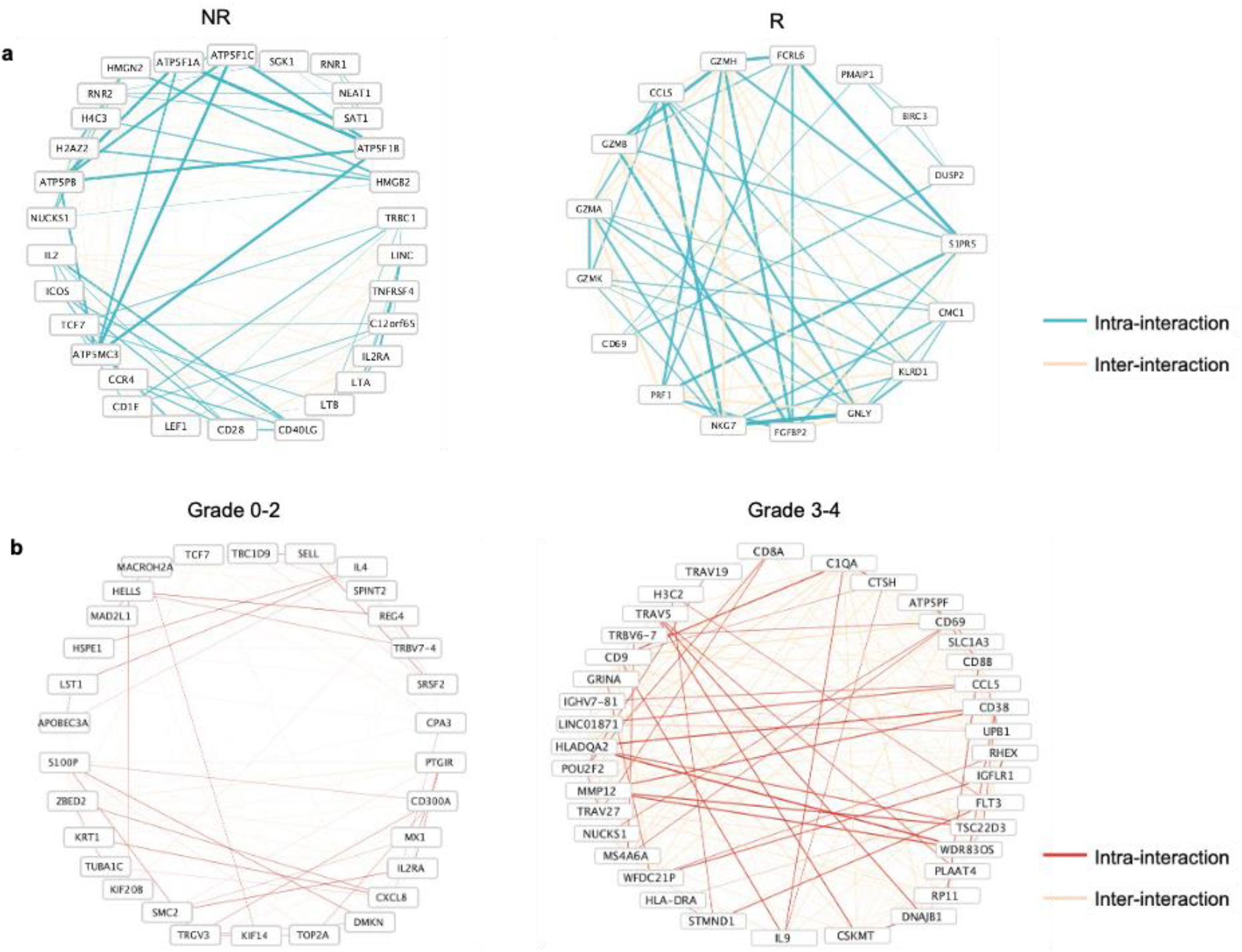
Gene regulatory networks inferred by CART-GPT. a, Regulatory networks (circle layout) for non-responders (left) and responders (right). Nodes represent high-contribution genes, and edges denote intra- and inter-metagene. Edge thickness indicates interaction strength. b, Regulatory networks (circle layout) for low-grade ICANS (left) and high-grade ICANS (right), with node and edge representations as in panel a.

**Extended Data Figure 9.**
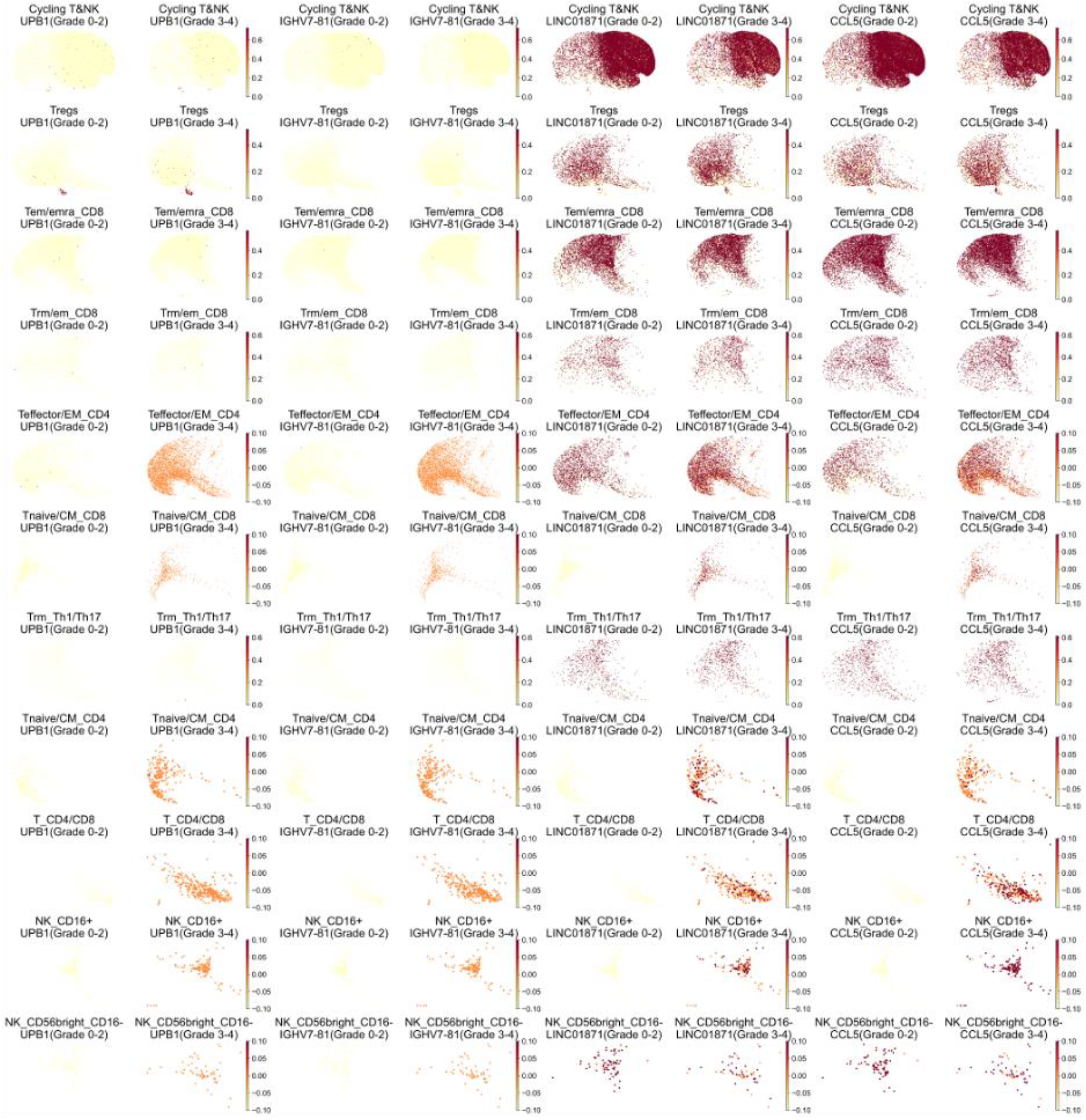
Cell-type–resolved expression of representative genes from the 65_SCORE metagene. Color intensity indicates gene expression, with darker shades reflecting higher expression. Each row represents a distinct cell subtype, and each column corresponds to a gene.

**Extended Data Figure 10.**
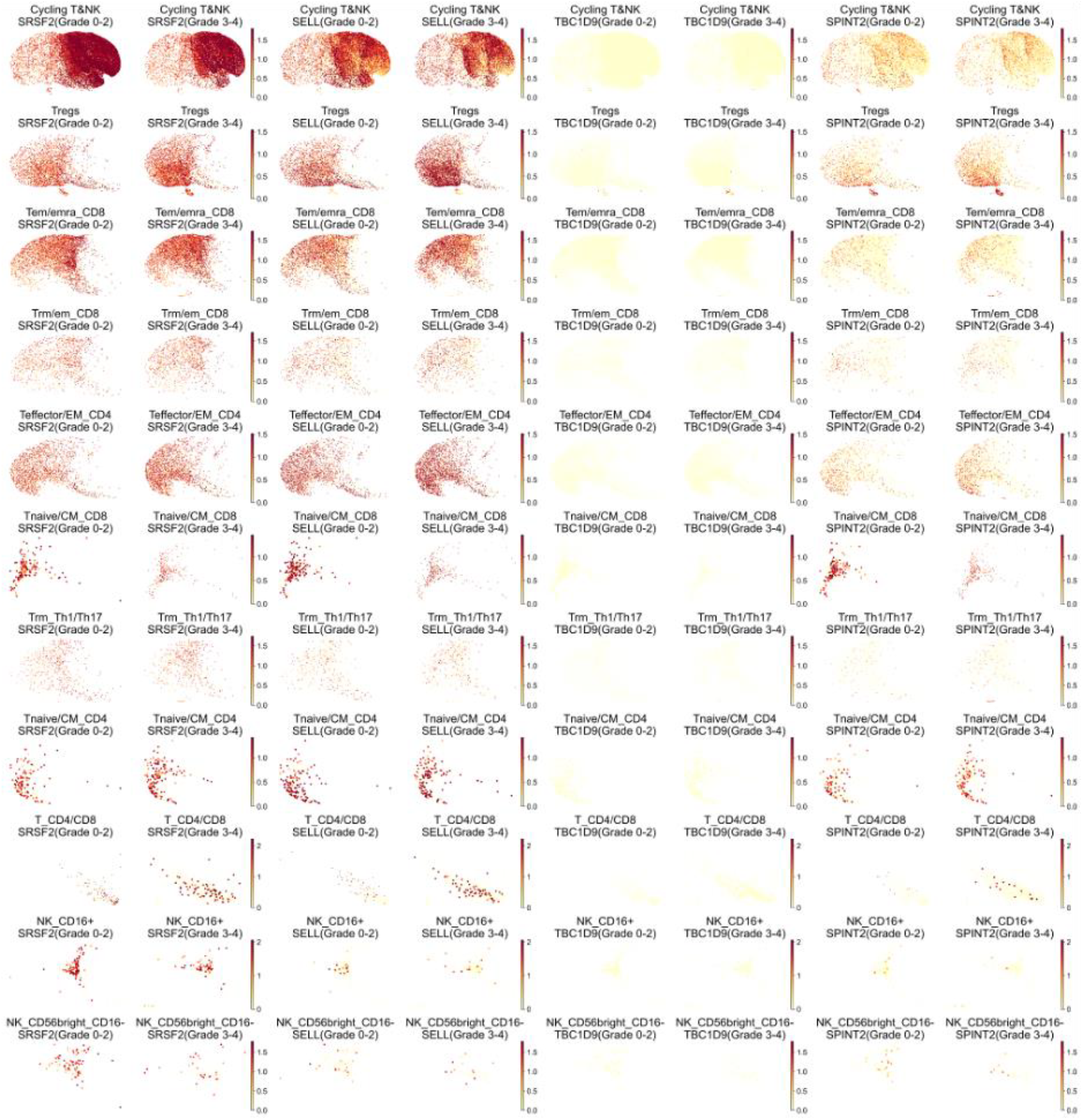
Cell-type–resolved expression of representative genes from the 35_SCORE metagene. Color intensity indicates gene expression, with darker shades reflecting higher expression. Each row represents a distinct cell subtype, and each column corresponds to a gene.

